# A planar dimer of bovine ATP synthase

**DOI:** 10.1101/2025.07.21.665874

**Authors:** Chimari Jiko, Atsuki Nakano, Yosuke Teshirogi, Eiki Yamashita, Genji Kurisu, Daron Standley, Tohru Terada, Kaoru Mitsuoka, Ken Yokoyama, Christoph Gerle

**Author notes:** These authors contributed equally (Chimari Jiko, Atsuki Nakano, Yosuke Teshirogi). Correspondence to Ken Yokoyama and Christoph Gerle.

## Abstract

Mammalian mitochondrial ATP synthase typically organizes into rows of V-shaped dimers that impose significant membrane curvature essential for mitochondrial cristae formation. Using gentle, column-free purification combined with single-particle cryo-electron microscopy, we have identified a previously unrecognized planar dimeric form of bovine ATP synthase exhibiting minimal membrane bending. This planar dimer is characterized structurally by antiparallel arrangement of two ATP synthase complexes linked by a straight conformation of the inhibitory factor 1 (IF1), a sharp contrast to the kinked IF1 observed in tetrameric assemblies. Molecular dynamics simulations confirm that transitioning between straight and kinked IF1 conformations occurs without significant energetic barriers. The planar dimer also displays distinct peripheral stalk positioning relative to its adjacent α subunit. These structural divergences suggest a specialized functional role and localization distinct from the canonical sharply membrane bending ATP synthase oligomers, providing structural support for a model of membrane curvature-driven division of labor within mitochondrial ATP synthase populations.

## Introduction

Mitochondrial F-ATP synthase/hydrolase, hereafter termed ATP synthase, plays a central role in biological energy conversion by catalyzing the interconversion between proton motive force (pmf) and the synthesis of adenosine tri-phosphate (ATP) through a rotary mechanism(1, 2). In all mitochondria examined thus far ATP synthase forms oligomers that induce positive membrane curvature by species dependent subunits that are not directly involved in rotary catalysis(3, 4). In mammalian mitochondria, ATP synthase forms long rows of V-shaped dimers that induce a strong ∼90° bend of the inner mitochondrial membrane at cristae ridges(5, 6). As a result, oligomers of ATP synthase, in conjunction with oligomers of the mitochondrial fusion protein Optic atrophy 1 (OPA1) located at cristae junctions, shape the architecture of the inner mitochondrial membrane (IMM)(7). Oligomer formation of mammalian ATP synthase is dynamic, and binding of the dimeric natural inhibitory factor 1 (IF1) to the matrix exposed F_1_ domain exerts a stabilizing effect(8). Higher oligomers of V-shaped ATP synthase dimers in actively respiring mitochondria are thought to occupy relatively stable locations at the cristae ridges(9). Under apoptotic stimuli directed at mitochondria, these higher-order oligomers disassemble, causing drastic alterations in cristae architecture and increasing the frequency of F_1_ domains positioned closer together than the known spacing observed in V-shaped dimers(10). Recently, it has become clear that the pmf of the inner mitochondrial membrane is not uniform; individual cristae and the inner boundary membrane (IBM) experience different levels of transmembrane electric potential (ΔΨ_m_)(11). Additionally, even under physiological conditions a single mitochondrion can harbor distinct populations of ATP synthase complexes, some engaged in ATP synthesis and others in ATP hydrolysis(12, 13). Moreover, live-cell microscopy has demonstrated that mammalian mitochondria contain subpopulations of ATP synthase complexes exhibiting either quasi-immobile or highly mobile behavior. An increase in the mobile subpopulation was observed in cells with inhibited glycolysis, and a higher proportion of these mobile ATP synthase complexes localized to the mitochondrial inner boundary membrane (IBM)(9, 14). Energetic considerations regarding membrane bending by mitochondrial ATP synthase have been proposed as a driving force for oligomerization, supported by both *in silico* simulations and *in vitro* experiments(15, 16). These studies indicate that a higher oligomerization state and reduced mobility are characteristic features associated with the pronounced membrane bending exerted by V-shaped dimers. In light of the above, the existence of an ATP-hydrolyzing ATP synthase subpopulation with higher mobility suggests the possibility of an oligomeric form structurally distinct from the established V-shaped dimer oligomers. To investigate this possibility, we examined whether oligomers composed of V-shaped dimers represent the sole oligomeric form of mammalian ATP synthase. To this end we developed a mild column-free large-scale purification method aimed at isolating IF1 stabilized oligomers of bovine ATP synthase at a level of purity and yield that is sufficient for structural characterization by single particle cryo-EM(17). Consistent with previous observations(18), we identified the smallest oligomeric unit of the V-shaped dimer as an IF1-bound tetramer of bovine ATP synthase, characterized by pronounced membrane bending. Additionally, we identified an IF1-bound dimer of bovine ATP synthase that exhibits only mild membrane bending, a structural feature compatible with localization in planar regions of the inner mitochondrial membrane. Therefore, we designated this complex the “planar dimer”.

## Results

To identify alternative oligomeric forms of mammalian ATP synthase, we developed a novel approach for isolating bovine ATP synthase from heart muscle mitochondria(17) (Fig.1A, fig. S1). Since oligomeric mammalian ATP synthase complexes are known to be highly sensitive to chemical and mechanical stress(19), we reasoned that avoiding mechanically stressful chromatography steps, employing buffers that promote binding of the oligomer-stabilizing natural ATPase inhibitory factor 1 (IF1), and using the gentle, lipid-like detergent glyco-diosgenin (GDN)(20) could facilitate the isolation of previously unidentified oligomeric forms of mammalian ATP synthase. Adjustment of buffer composition and sucrose density gradient conditions allowed column-free isolation of solubilized, IF1 stabilized oligomeric ATP synthase in milligram amounts at a level of purity sufficient for structural analysis (fig. S1). Cryo-EM image processing of oligomeric bovine ATP synthase (fig. S2, Table S1&S2) revealed two predominant oligomeric forms, both abundant and stable enough to be structurally determined at alpha-helical resolution: an IF1-stabilized tetramer resolved to an overall resolution of 7.2 Å, and an IF1-stabilized planar dimer resolved to an overall resolution of 5.0 Å (Fig. 1B,C).

**Figure 1.**
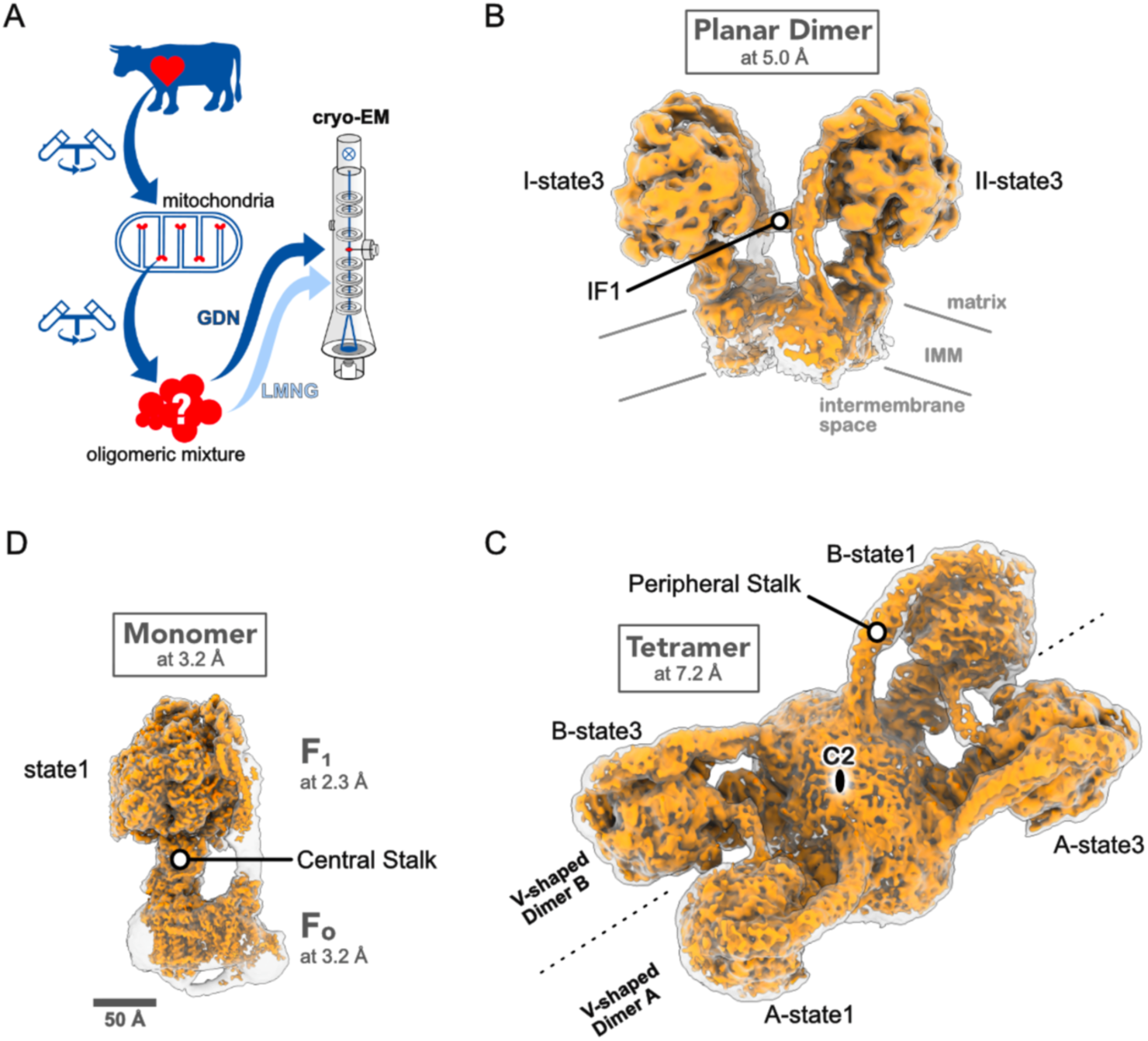
Oligomeric forms of bovine ATP synthase. **A** Workflow of ATP synthase preparation from bovine heart muscle tissue by differential and gradient centrifugation in the presence of the lipid like detergent GDN for preparation of oligomeric ATP synthase and LMNG for preparation of monomeric ATP synthase. **B** Cryo-EM map of the bovine planar dimer visualized at higher isosurface threshold (orange) and lower threshold (transparent). **C** Cryo-EM map of the bovine tetramer visualized at higher isosurface threshold (orange) and lower threshold (transparent). V-shaped dimer A and B are separated by a dashed line and the C2 symmetry center is indicated. **D** Cryo-EM map of the bovine monomer with the F_1_ focused map and the F_o_ focused map visualized in orange and the overall map in transparent. Scalebar in (D) applies also to (B) and (C).

### IF1 bound tetramer of bovine ATP synthase

In the IF1 bound bovine ATP synthase tetramer, two V-shaped dimers of the bovine ATP synthase are aligned side-by-side with two dimers of IF1 each interlinking two adjacent F_1_ domains (Fig.1C). The rotary states(21) of neighboring F_1_ domains interconnected by a single IF1 dimer are in state 1 and state 3, respectively (fig. S3). This arrangement of the four protomers results in a C2 symmetry for the tetramer, with each protomer denoted by being part of the V-shaped dimer A or B and its rotary state being 1 or 3: A-state1, A-state3, B-state1 and B-state3. The peripheral stalks (PS) of the IF1 interlinked neighboring F_1_ domains are on the same side, facing those of the opposite IF1 interlinked F_1_ domains. The membrane bending b-e-g unit of each protomer is close to the C2 symmetry center of the tetramer, which results in an arching convex architecture for the membrane spanning part of the whole tetramer, determining the local membrane curvature. Thus, the overall architecture of the bovine ATP synthase tetramer closely resembles that of the previously described porcine heart mitochondrial ATP synthase tetramer(18). Our 7.2 Å cryo-EM map-based main chain model of the bovine tetramer visualizes some of the protein-protein interactions that might play a role in the spatial organization of two V-shaped dimers (fig. S4). The clearest and most important interaction is mediated by the two dimers of IF1 that intercalate the neighboring F_1_ domains of each V-shaped dimer. The complete insertion of all four N-terminal regions results in equal distances between both IF1 binding sites—a configuration likely contributing to the overall C2 symmetry observed in this 2.4 MDa complex. Additional sites of protein-protein interaction between the two V-shaped dimers can be found both at the surface and the interior of the membrane spanning F_o_ domain. Although the current reconstruction does not permit definitive conclusions, we wish to highlight three potential interaction sites between the two V-shaped dimers mediated by subunits DAPIT (termed subunit k in yeast), g, and e (fig. S4). Subunit g appears to establish contacts both with itself and with DAPIT at the membrane surface, while the C-terminus of DAPIT may also form an intramembrane interaction with the helix 5–helix 6 turn of subunit a. In addition, opposing subunits e of the A-state1 and B-state1 protomers appear to form an intramembrane contact. The C-terminus of subunit e contacts the luminal density of the c-ring protruding into the intercristae space, and apparently not any of the c subunits itself, which is an important observation that was predicted(22) and previously reported for the porcine tetramer, the bovine monomer and dimer as well as for ovine and human monomers(18, 21, 23, 24, 25, 26). This particular contact allows free Brownian rotation of the c-ring, an essential prerequisite for proton driven rotary catalysis. In the bovine tetramer the C-terminal regions of all four e subunits are in proximity, possibly forming an additional intra V-shaped dimer contact. We note, however, that to draw firm conclusions regarding intramembrane interactions between V-shaped dimers, all the potential contacts within the F_o_ domains of the ATP synthase tetramer described above must be further characterized using maps of significantly higher resolution than currently available.

### IF1 bound planar dimer of bovine ATP synthase

The structure of the IF1 bound planar dimer is, to our knowledge, a previously undescribed dimeric form of bovine ATP synthase. It features two adjacent ATP synthase complexes arranged side-by-side in an antiparallel orientation and interconnected by a single, straight IF1 dimer (Fig. 2). In contrast to the architecture of the tetramer, this results in the peripheral stalks (PS) of the two ATP synthase complexes being on opposite sides of the interlinking IF1 dimer. Both ATP synthase complexes in the planar dimer exhibit density for almost all known subunits, for which we constructed a main-chain model using PDB ID 6ZIU and 6Z1U(22) as the initial reference structures. Of note, the density corresponding to subunit 6.8PL appears to be absent, whereas cryo-EM density indicative of DAPIT is present (fig. S5). Both subunits are believed to be directly involved in oligomerization and thus their presence or absence is of interest. However, since the overall transmembrane density for the planar dimer is weak, we chose to exclude them from our model. A confident judgement on the presence of 6.8PL and DAPIT in the planar dimer and their potential role in switching between V-shaped dimer to planar dimer will have to await further studies. Clear density for the IF1 dimer that interlinks the two neighboring F_1_ domains allowed the fitting of a main chain model of its X-ray crystal structure (PDB ID 1GMJ)(27). Likewise, the well resolved density for the catalytic α_3_β_3_ hexamer, its central rotor γ subunit and the peripheral stalk clearly showed that both complexes are in the same rotary state: state 3 (fig. S3).

**Figure 2.**
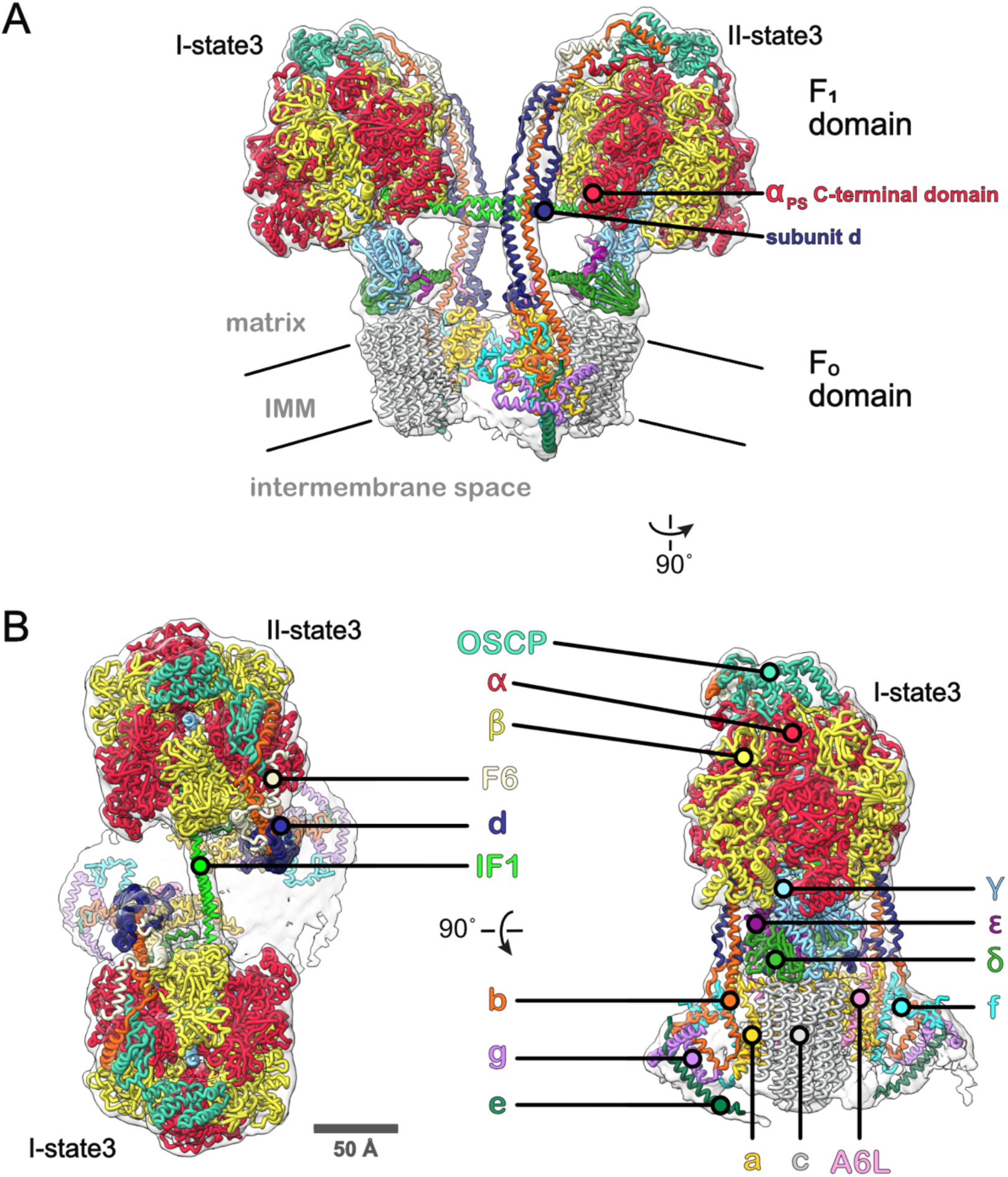
Structure of the bovine planar dimer. **A** Frontal side view along the membrane plane with the cryo-EM map depicted in transparent silhouette style and the main chain model of the subunits in thick wire style and colored as their name font in **B**. **B** Left panel: ‘top’ view of the planar dimer from the matrix side with map and model as in **A**. Right panel: side view along the long axis of the planar dimer with map and model as in **A**.

### Membrane bending architecture of oligomeric bovine F-ATP synthase

As its name suggests, the most distinctive feature of the planar dimer is its compatibility with localization in comparatively planar regions of the inner mitochondrial membrane. These regions could include the inner boundary membrane (IBM) or the extended, planar portions of cristae that project into the matrix. In contrast, the well-characterized V-shaped dimer of bovine ATP synthase induces substantial membrane curvature: the b–e–g subunit interface creates ∼45° of bending per monomer, resulting in an overall ∼90° bend when dimers are arranged in extended rows(23). The IF1-stabilized tetramer described here further illustrates this curvature by visualizing an oligomer of two V-shaped dimers (Fig.3B). To estimate membrane bending, we used the relative position of the c-rings as a structural proxy—based on the observation that isolated c-ring subcomplexes form flat 2D crystals in reconstituted lipid bilayers and thus do not intrinsically bend membranes (28). In the bovine tetramer, the rotor rings span a maximal angle of ∼115°, indicating a more pronounced curvature than previously seen in cryo-electron tomography of mitochondria(5) or in single-particle cryo-EM of purified dimers(29). This increased bending mirrors prior findings from the IF1-bound porcine ATP synthase tetramer(18). In contrast, the planar dimer exhibits a c-ring angle of just ∼35°—dramatically less than both the V-shaped dimer (∼90°) and the ∼45° curvature associated with monomeric ATP synthase reconstituted in lipid bilayers(19). These geometric differences suggest that the planar dimer imposes minimal energetic costs when embedded in flat membrane regions, distinguishing it from the highly curvature-inducing V-shaped dimers.

**Figure 3.**
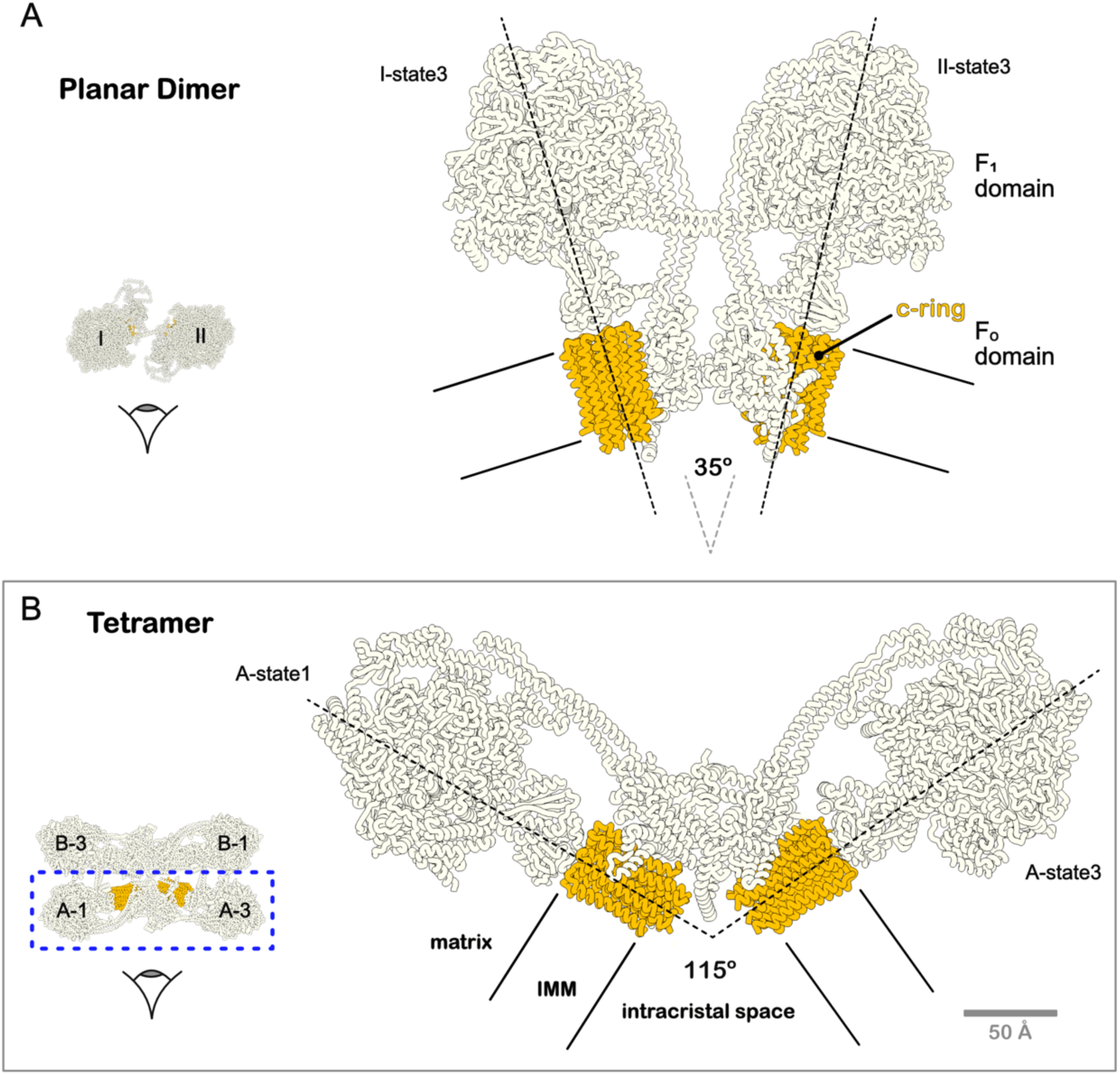
Membrane bending of oligomeric bovine ATP synthase. **A** Frontal side view of the planar dimer in thick wire style with the c-ring depicted in orange and all other subunits in white. The long axis of each protomer is indicated by long dashed lines and their relative angle of 35° indicated between them. The assumed position of the inner mitochondrial membrane is indicated by solid lines. Viewing direction as indicated by the left side inset. **B** Frontal side view of tetramer in thick wire style with the c-ring depicted in orange and all other subunits in white. The long axis of each protomer is indicated by long dashed lines and their relative angle of 115° indicated between them. The assumed position of the inner mitochondrial membrane is indicated by solid lines. Viewing direction as indicated by the left side inset.

### Conformations of IF1

Consistent with our experimental design (fig.S1B), dimeric IF1 is firmly bound to both the planar dimer and the ATP synthase tetramer (Fig. 4A,B). Clear density for the IF1 dimer enabled main-chain model fitting and suggests full occupancy, comparable to that of the α₃β₃ catalytic hexamer. As in the previously reported IF1-bound porcine tetramer(18), the IF1 dimer intercalated between neighboring F1 domains adopts a markedly kinked conformation—unlike the straight structure seen in the X-ray crystallography of isolated IF1 dimers(27) (Fig. 4B,E). Specifically, the N-terminal α-helices bend sharply after the dimerization domain, turning to insert into the α–β interface of the adjacent F1. In the B-state3 protomer, this bend measures ∼56°, highlighting the conformational flexibility between the N-terminal and C-terminal regions. Our structure confirms this kinked IF1 conformation in the bovine ATP synthase tetramer. In stark contrast, the IF1 dimer linking the two F1 domains of the planar dimer adopts a straight, unbent conformation (Fig. 4A).

**Figure 4.**
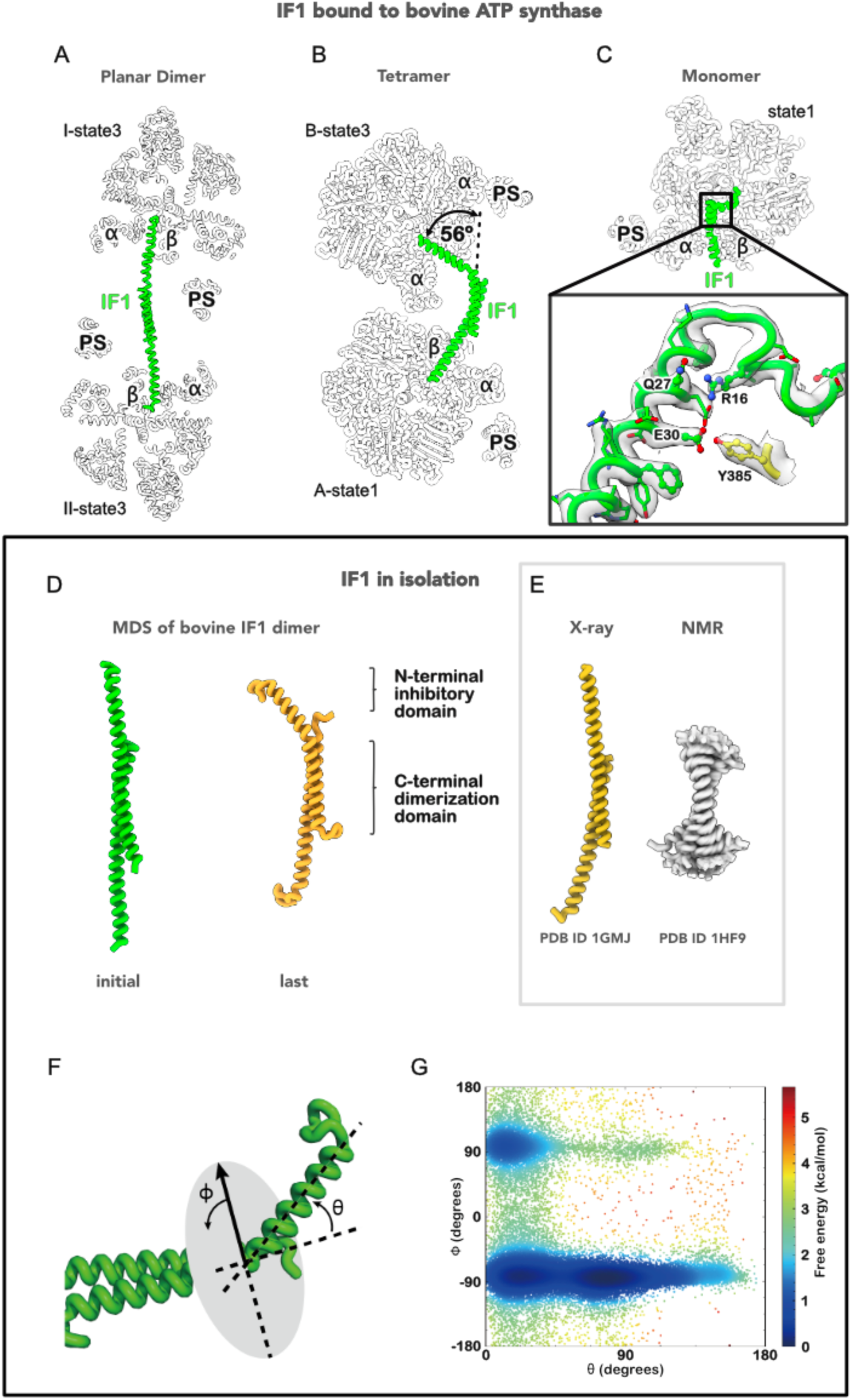
Conformations of IF1. **A** Cross-sectional view of the planar dimer parallel to the IF1 dimer. The almost straight, two neighboring F1 intercalating IF1 dimer is depicted in green and all other subunits in white. **B** Cross-sectional view of the tetramer A-state1 and B-state3 F1 domains parallel to the IF1 dimer. The strongly kinked, two neighboring F1 intercalating IF1 dimer is depicted in green and all other subunits in white. **C** Cross-sectional view of IF1 bound state 1 monomer F1 domain. The inset shows map and model of the inhibitory N-terminal region of IF1. High-lighted are the self-interaction between glutamine 27 and arginine and the interaction between glutamate 30 of IF1 and tyrosine 385 of subunit β. **D** The initial (left, green) and the last (right orange) structures of the 500-ns MD simulation conducted for the symmetric model of bovine IF1 dimer. The ranges of the N-terminal inhibitory domain and the C-terminal dimerization domain are indicated. **F** Definition of θ and φ angles describing the orientation of bending of the N-terminal inhibitory domain with respect to the C-terminal dimerization domain. **G** Distribution of the angles of IF1 in the MD simulation conducted for the symmetric model. Each point represents an individual bending angle and is colored by the free-energy value.

This observation raises the question of whether the kinked and straight conformations of the IF1 dimer in oligomeric ATP synthases represent energetically distinct states—specifically, whether transitioning between these conformations requires external energy input. To obtain insights into this question, we constructed a symmetric IF1 dimer *in silico* model and performed molecular dynamics (MD) simulation. In agreement with the expectation that the two IF1 monomers formed a stable binding interface with one another at the C-terminus, during the time of simulation the α-helical structure of the IF1 dimer was maintained throughout the simulation (Figure 4D). Meanwhile, frequent bending was observed in the N-terminal domain, where the residues exhibited less interaction between the IF1 monomers (Figure 4F; Supp. movie md1). Furthermore, we observed bent IF1 dimer conformations closely resembling those determined by cryo-EM for the ATP synthase tetramer-bound state (fig. S6). This bending behavior was consistently reproduced across simulations, including those incorporating post-translational modification (succinylation of Lys78) as well as IF1 dimers from other species, such as porcine and human (Supp. Movies md2–4). Interestingly, the succinylation of lysine 78 resulted in salt bridges between His49 and Lys46 in the same monomer that stabilized the bent conformation of the IF1 dimer (fig. S7), suggesting a potential mechanism for bending. Despite this interaction, each IF1 monomer retained flexibility, as evidenced by the absence of a significant free-energy barrier (<0.6 kcal/mol ≈ *kT*) associated with the θ-angle describing the relative orientation of the N-terminal domain (Figure 4F). Taken together, the MD simulations of the IF1 dimer in the absence of other proteins support the notion that the kinked and non-kinked conformations of the IF1 dimer bound to oligomeric ATP synthase do not require any external energy input greater than *kT* for their conformational interchange at ambient temperatures.

For more detailed analysis of how physiologically bound IF1 interacts with the F1 domain, ATP synthase oligomers were monomerized by substituting the relatively gentle detergent GDN with the harsher amphiphile lauryl maltose-neopentyl glycol (LMNG)(30) in the final sucrose density gradient. Structure determination by single particle cryo-EM yielded a map of the state 1 monomer with the F_1_ domain resolved at a resolution of 2.3 Å, enabling us to obtain detailed direct structural insights into how physiologically bound IF1 interacts with the F_1_ domain (Fig.4C). The observed interactions were qualitatively identical to those observed in the high resolution (2.1 Å) X-ray crystal structure of the isolated bovine F_1_ domain bound with the monomeric N-terminal fragment of IF1 termed I1-60His (PDB ID 2V7Q)(31). These included hydrophilic interactions between IF1 and the β subunit, such as β tyrosine 385 with IF1 glutamate 30 (Fig.4C inset), or between IF1 and the central stalk (CS) γ subunit, such as γ asparagine 15 with IF1 serine 10 and γ isoleucine 16 with IF1 phenyl alanine 22 (fig. S8), interactions crucial for mediating directional inhibition in the ATP hydrolysis direction(32). A notable deviation from these matching interactions is the differing conformation of IF1 arginine 16 from that seen in the X-ray crystal structure. In our cryo-EM structure the side chain of IF1 arginine 16 is in close contact to IF1 glutamine 27 (3.2 Å from the Nη of the Arg16 guanidino group to the Oγ of the Gln27 amide group) (Fig.4C inset). This side chain arrangement is likely to form a hydrogen bond, a notion that is supported by a map-to-map contact visualized at a threshold that also visualizes the contact between βTyr385 and IF1Glu30. This self-interaction is also apparent in the single particle cryo-EM structure of the bovine ATP synthase dimer (PDB ID 6ZQN)(23), possibly indicating a slight difference for IF1 binding to the isolated F_1_ domain *vs* the intact ATP synthase complex.

### Proximity of PS and α_PS_

A distinct structural feature of the planar dimer is the unusual position of the peripheral stalk relative to the C-terminal domain of the α subunit that is binding to the PS, here termed α_PS_. In mammalian ATP synthase the peripheral stalk is, in comparison to its bacterial or chloroplast counterpart, an elaborate structural element comprising subunit OSCP, b, F6, d and parts of A6L. It exhibits a left-handed curved overall structure in the resting state, i.e., absence of any driving force. Subunit α_PS_ of the F_1_ domain forms an intricate contact with OSCP, subunit F6 and subunit b *via* its short N-terminal alpha helix, while its C-terminal region in the absence of a driving force is in close proximity to subunit d. This appears to be the case for all single particle cryo-EM structures of monomeric mammalian ATP synthase reported to date(21, 23–26). In contrast, for both ATP synthase complexes of the planar dimer the peripheral stalk is clearly separated from the C-terminal domain of the α_PS_ subunit, leaving a gap of more than 15 Å between the closest main chain C_α_ atoms of subunit d and the C-terminal α_PS_ (Fig.5A). In the ATP synthase tetramer, the situation is more complex: peripheral stalks of ATP synthase protomers in rotary state 1 adopt a conformation resembling that observed in the planar dimer, whereas peripheral stalks of protomers in rotary state 3 assume conformations more akin to monomeric bovine F-ATP synthase, characterized by close proximity between subunit d of the peripheral stalk and the C-terminal domain of α_PS_ (Fig. 5B,C).

**Figure 5.**
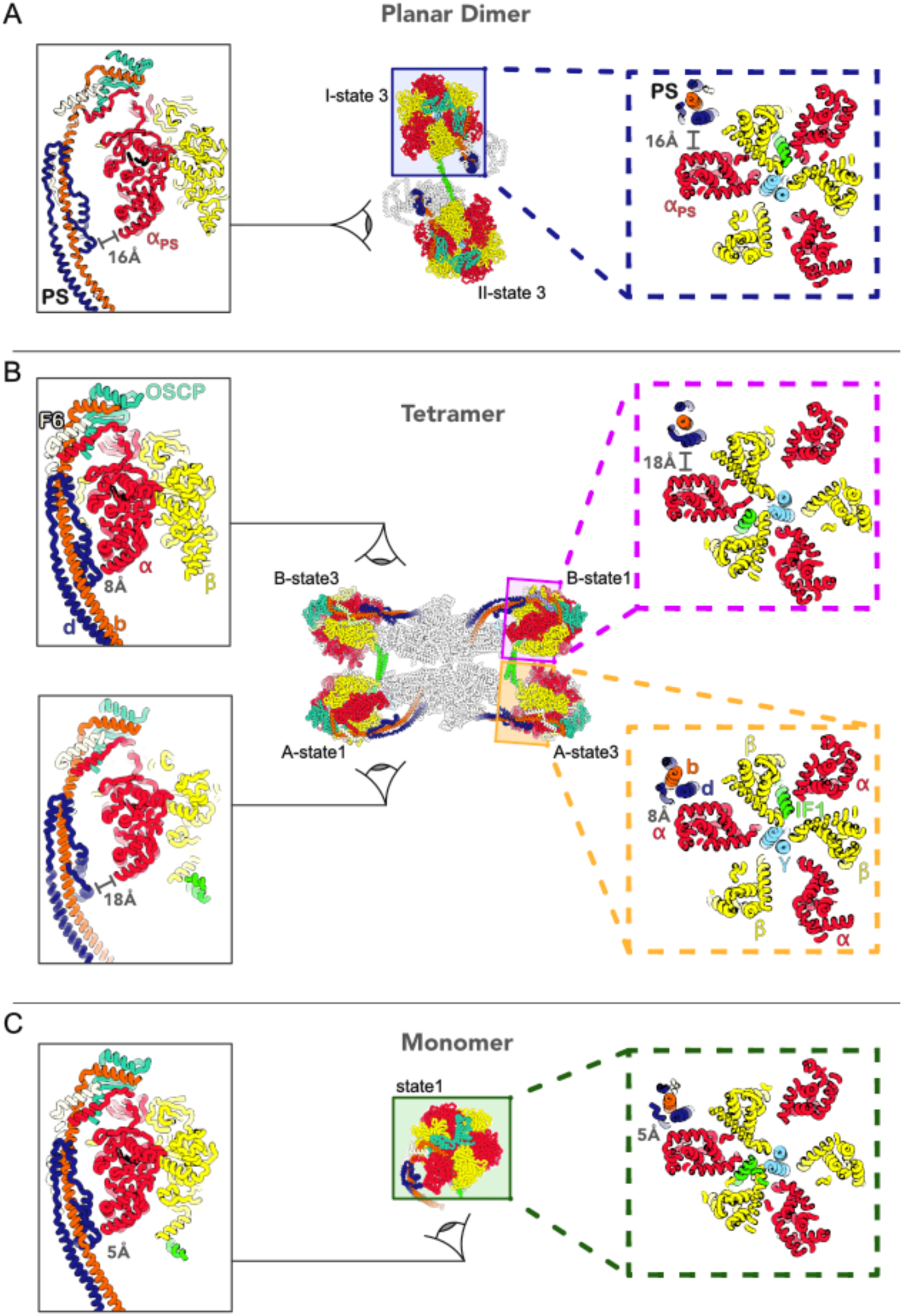
Proximity of Peripheral Stalk and C-terminal α_PS_. **A** Left panel: Side view of the planar dimer II-state3 protomer showing the peripheral stalk and α_PS_. The gap of 16 Å at the narrowest point between C_α_ atoms of subunit d and α_PS_ is indicated. Right panel: Cross-sectional view of the planar dimer I-state3 protomer F_1_ domain at the height of α_PS_ C-terminal region. The gap of 16 Å at the narrowest point between C_α_ atoms of subunit d and α_PS_ is indicated. Subunit coloring as in Figure 2. **B** Upper left panel: Side view of the tetramer B-state3 protomer showing the peripheral stalk and α_PS_. The gap of 8 Å at the narrowest point between C_α_ atoms of subunit d and α_PS_ is indicated. Lower left panel: Side view of the tetramer A-state1 protomer showing the peripheral stalk and α_PS_. The gap of 18 Å at the narrowest point between C_α_ atoms of subunit d and α_PS_ is indicated. Upper right panel: Cross-sectional view of the tetramer B-state1 protomer F_1_ domain at the height of α_PS_ C-terminal region. The gap of 18 Å at the narrowest point between C_α_ atoms of subunit d and α_PS_ is indicated. Lower right panel: Cross-sectional view of the tetramer A-state3 protomer F_1_ domain at the height of α_PS_ C-terminal region. The gap of 8 Å at the narrowest point between C_α_ atoms of subunit d and α_PS_ is indicated. Subunit coloring for the F_1_ domain as in Figure 2. **C** Side view of the monomer state 1 showing the peripheral stalk and α_PS_. The gap of 5 Å at the narrowest point between C_α_ atoms of subunit d and α_PS_ is indicated. Right panel: Cross-sectional view of the F_1_ domain at the height of α_PS_ C-terminal region. The gap of 5 Å at the narrowest point between C_α_ atoms of subunit d and α_PS_ is indicated. Subunit coloring as in Figure 2.

## Discussion

Here, we investigated the oligomeric state and structure of bovine F-ATP synthase from heart muscle mitochondria by combining very mild column-free purification procedures that stabilize IF1 binding with single particle cryo-EM. Our approach enabled structural visualization of both the sharply membrane-bending, IF1-bound ATP synthase tetramer and the mildly bending, IF1-bound planar dimer. Throughout the purification process—beginning with the homogenization of freshly isolated heart muscle tissue—we used only buffers that promote IF1 binding, while deliberately avoiding the addition of exogenous ATP or IF1(17). These experimental circumstances give us confidence that the IF1 dimer present in the structures of planar dimer and tetramer did bind to the ATP synthase at the time of tissue collection.

An IF1 bound ATP synthase tetramer consisting of two V-shaped dimers connected by two IF1 dimers that intercalate neighboring F_1_ domains had previously been described by single particle cryo-EM of crude extracts from the inner mitochondrial membrane isolated from pig heart muscle tissue at ∼6 Å(18). A similar structure, albeit at the considerably lower resolution of ∼20 Å, was also found in isolated ATP synthase fractions from sheep heart muscle tissue(24). Furthermore, studies by cryo electron tomography of the inner membrane of mammalian mitochondria clearly showed that *in situ* ATP synthase is organized as long rows of membrane bending V-shaped dimers at the ridge of cristae membranes(5). These previous findings reinforce the notion that the bovine tetramer structure described here reflects the oligomeric state of ATP synthase found in the inner mitochondrial membrane before tissue homogenization.

What could be the possible location of the planar dimer in the inner mitochondrial membrane? Its membrane-bending properties render it viable in regions of either slight positive curvature or even close to planar regions of the inner boundary membrane. Therefore, we propose that the planar dimer represent the highly mobile subpopulation of ATP synthase complexes that are mainly engaged in proton pumping powered by ATP hydrolysis located at the inner boundary membrane (IBM)(9). Under physiological conditions, this mechanism is thought to help sustain the proton motive force needed for mitochondrial transport; however, in pathophysiological contexts, it may contribute to ATP depletion and energy failure(33). A high level of mobility should also exist for ATP synthase complexes that are not fully assembled yet and are *en route* to higher oligomer formation. However, the cryo-EM density for subunit a of the planar dimer in our structure is strong, clearly indicating its stoichiometric presence in the planar dimer. And since subunit a is added in the last step of complex assembly(34), the interpretation of the planar dimer as an assembly intermediate can be ruled out.

Apart from its only mildly membrane bending nature, a distinct structural feature of the planar dimer is the clear detachment of subunit d from the C-terminal region of subunit α_PS_. This is remarkable, because the proximity of the peripheral stalk to the C-terminal region of α_PS_ is a conserved structural feature of ATP synthase from bacteria, chloroplasts, and mitochondria. Furthermore, for bacterial ATP synthase this attachment is seemingly steady even during catalytic turnover(35). An exception to this widespread structural motif in F-type ATP synthases are the peripheral stalks in unicellular algae *Polytomella* sp. and *Chlamydomonas reinhardtii*, where this attachment is clearly absent(36, 37). And more notable, clear detachment of the PS from α_PS_ has been visualized for yeast ATP synthase under the strain from ATP hydrolysis powered rotation(38). The widespread occurrence of physical interactions between the peripheral stalk and the C-terminal region of α_PS_ suggests a potential functional advantage for rotary catalytis. Additionally, the clear detachment of the peripheral stalk subunit d from α_PS_ in both planar dimer and two protomers of the tetramer point to the possibility that a reversible detachment of the peripheral stalk is a specific function of subunit d.

The IF1 dimer that connects the two F_1_ domains of the planar dimer is straight, whereas in the bovine tetramer, in confirmation of what has been reported for the porcine tetramer, it is kinked between the C-terminal dimerization domain and the inhibitory N-terminal domain (Fig.4). Our molecular dynamics simulations suggest that no energy barrier greater than *kT* prevents the interconversion of straight and kinked conformations for the physiologically bound IF1 dimer. Thus, mammalian IF1 appears to have evolved into an inhibitory protein that is equally able to bind and stabilize two structurally very distinct oligomeric forms of mitochondrial ATP synthase. The IF1 bound to the bovine ATP synthase reported here is the physiological full length dimer form present in heart muscle tissue mitochondria and the high-resolution structure of the F_1_ intercalating inhibitory N-terminal domain is identical to that analyzed by X-ray crystallography of the heterologous expressed and His-tagged N-terminal fragment of monomeric IF1, consisting of residues 1-60 plus a C-terminal His_6_ *in vitro* bound to the isolated F_1_ subcomplex(31). In this respect, our high-resolution structure confirms the relevance of the *in vitro* data obtained in experimental set-ups, such as single molecule rotation assays, that use the isolated F_1_ subcomplex inhibited by N-terminal fragments of IF1(32). Previously reported 2D and 3D negative stain EM and cryo-EM image data of mitochondrial ATP synthase at low resolution appear to be obtained from the same oligomeric entity as the planar dimer(10, 23, 25, 39–41)(fig. S9). Therefore, we believe that the planar dimer is not rare, had been isolated before and was simply overlooked due to the lack of unambiguous data. In the last few years cryo-EM has revealed a surprising diversity of oligomeric forms of F-ATP synthase/hydrolase present in mitochondria from various organisms(4). Our study now demonstrates that for mammalian mitochondria two different oligomeric forms of ATP synthase co-exist, implicating a membrane-curvature driven spatial and functional division of labor for this central energy converter of the mammalian cell.

## Methods

### Purification of bovine ATP synthase

The column-free purification of IF1 bound bovine ATP synthase from bovine heart muscle tissue was performed as previously described in detail (Fig. S1) (17). In brief, the freshly slaughtered cow’s heart tissue was minced, homogenized and mitochondrial inner membranes were collected by centrifugation. Inner membranes were homogenized using a Dounce glass homogenizer in a buffer of 40 mM HEPES-Na (pH 7.3), 5 mM EGTA, 5 mM DTT, 5 mM MgCl_2_ and 0.5 mM ADP, solubilized by the addition of deoxycholic acid (0.73%), decyl-maltoside (0.4 %) and glyco-diosgenin (GDN) (0.1%) and subjected to a step/equilibrium (2.3 M and 1.6 M) sucrose density ultracentrifugation for 42 hours at 176,000*g* using a buffer containing 40 mM HEPES-Na (pH 7.3), 5 mM DTT, 5 mM EGTA, 5 mM MgCl₂, 0.5 mM ADP, 100 mM KCl, 0.1% DM, 0.02% LMNG and 0.02% GDN. Resulting fractions were examined by SDS-PAGE, CN-PAGE, negative stain EM and ATPase activity measurement for the presence of ATP hydrolase activity free oligomeric ATP synthase complexes and pooled. A second step (2.3 M, 1.6 M, 1.0 M and 0.6 M) sucrose density gradient at 176,000*g* was conducted for 20 hours using a buffer of 40 mM HEPES-Na (pH 7.3), 5 mM DTT, 5 mM EGTA, 5 mM MgCl₂, 0.5 mM ADP, 0.02% GDN, and 80 mM KCl and the resulting fractions examined by SDS-PAGE, CN-PAGE, negative stain EM and ATPase activity measurement for the presence of ATP hydrolase activity free oligomeric ATP synthase complexes and pooled for a final linear (20 - 40%) sucrose density gradient ultracentrifugation at 113,000×g for 20 hours using a buffer containing 40 mM HEPES-NaOH (pH 7.3), 5 mM MgCl_2_, 5 mM DTT, 5 mM EGTA, 0.5 mM ADP, 80 mM KCl, and 0.02% GDN. For the preparation of monomeric ATP synthase GDN was replaced with LMNG (0.5%). Resulting fractions enriched in either oligomeric or monomeric ATP synthase were pooled and used for cryo-grid preparation.

### CryoEM grid preparation

Prior to cryo-grid preparation sucrose was removed from the sample buffer via PEG-20,000 precipitation and the concentration adjusted to ∼0.5 mg/ml. For all data sets cryo-grids were prepared using Quantifoil R2/2 Cu + 2nm continuous carbon grids that had been freshly treated to 30 sec glow discharge using a Plasma Ion Bombarder Vacuum Device (PIB-10). ATP synthase solution (3.5 μl) was applied to the cryo-grid and blotted for 0.5-1 sec (blot force 10, wait time 10s, drain time 0.5 sec at 100% humidity and 4°C) using a FEI Vitrobot Mark IV (ThermoFisher), followed by flash freezing in liquid ethane. The prepared cryo-grids were stored in liquid nitrogen until use.

### Cryo-EM data acquisition

Cryo-EM imaging was performed using a Titan Krios (FEI/Thermo Fisher Scientific) operating at 300 kV acceleration voltage and equipped with a K3 electron detector (Gatan) in electron counting mode (CDS). SerialEM(42) was used for data collection. Movies at 50 frames each were collected at a nominal magnification of 88,000 with a pixel size of 0.88 Å/pix using a defocus range of 0.8-2.0 μm and employing a total electron dose of 50 electrons/Å_2_.

### Image processing

#### Tetramer

The detailed workflow for single-particle analysis is summarized in Supplementary Figure 2. RELION(43) and CryoSPARC(44) were used for image analysis. File format conversion from CryoSPARC to RELION was performed using csparc2star.py from pyem., beam-induced drift was corrected for using MotionCor2(45), and CTF estimation was performed using CTFFIND4(46). Density maps for the initial model of the tetramer were generated from 1,000 micrographs. Particles were first manually picked for training a topaz(47) model. They were then extracted with a box size of 200 pixel (3.36Å/pixel) and 2D classification was performed. Particles from good 2D class average images were selected and a topaz model was further trained to extract the picked particles. Particles in the 2D class-averaged image that appeared to be ATP synthase tetramers were selected for a multi-class ab-initio reconstruction. An initial map of the tetramer was obtained by 3D classification in RELION followed by 3D refinement with C2 symmetry imposed. As in the initial model creation, 5,734,709 particles were picked and extracted by topaz from all 48,384 micrographs. Multiple 2D classifications selected 1,277,913 good particles, and 3D classification was performed. The particles were then re-extracted at 600pix (0.84 Å/pixel) and 3D refinement was performed with C2 symmetry imposed. We performed symmetry expansion based on C2 symmetry and carried out focused refinement for each of the two unique F_o_F_1_ complexes individually. Subsequently, Bayesian polishing and CTF refinement were applied to improve the resolution. Symmetry expansion based on C2 symmetry was performed, followed by focused refinement of each individual F_o_F_₁_ complex and a further round of Bayesian polishing and CTF refinement. Using the volume eraser in ChimeraX(48), the respective densities of F_1_ and F_o_ were erased from the obtained density map of F_o_F_1_, and the masks of F_1_ and F_o_ were created with RELION. Using this mask, the 3D structures of F_1_ and F_o_ were reconstructed by focused refinement. The structure of the entire ATP synthase tetramer and the central region was reconstructed by 3D refinement, using the CTF parameters optimized in the F_1_ part, after removing duplicate particles by symmetry expansion.

#### Planar dimer

Autopicking was performed on 21,772 preprocessed micrographs using a Topaz model trained on a tetramer dataset. The extracted particles with a 200 pixel box size (3.36Å/pix) were subjected to multiple 2D classifications, and 34,002 particles corresponding to 2D class averages that appeared to represent planar dimers were selected. Ab-initio Reconstruction was performed in five classes, yielding initial maps that appeared to be monomers, V-shaped dimers, and planar dimers. Tetramer structures were added to the five maps obtained by ab-initio reconstruction, and heterogeneous refinement was performed on 290,036 particles selected by 2D classification. Planar dimer structures reconstructed from 136,954 particles were further classified by heterogeneous refinement based on the relative positions between monomers of planar dimers. After re-extraction of the particles with a 250 pixel box size (2.18Å/pixel), Bayesian polishing was performed with RELION. The 3D structure was reconstructed at 5 Å by local refinement using planar dimer masks.

#### Monomer

Initially, particles were manually picked from 1,000 micrographs and used to train a Topaz model for automated particle picking. The picked particles were subjected to 2D classification, and 2,000 representative particles were selected to further train the Topaz model. The retrained model was then applied to 19,983 micrographs for particle picking. A total of 5,605,840 particles were extracted with 90 pixel box size (3.36Å/pixel) and subjected to 2D classification, resulting in the selection of 1,917,221 particles. An initial model was created with ab-initio reconstruction using 100,000 of those particles, and heterogeneous refinement was performed iteratively. Heterogeneous refinement resulted in a structure consisting of 398,620 particles in rotary state1, 117,760 in rotary state2, and 488,235 in rotary state3. Each class was re-extracted at 240 pixel box size (1.26 Å/pix) and 3D refinement was performed. The resolution was improved by CTF refinement and Bayesian polishing after local refinement of the F_1_ domain. Rescaling to 340 pixel box size (0.94Å/pixel) was also performed during the Bayesian polishing. In State 3, where the resolution was most improved, local refinement was performed by a mask containing F_o_ and γ. Then, 3D classification was performed without alignment in RELION to select the classes with the best resolution.

### Model building

The bovine IF1 bound tetramer model was built using the bovine ATP synthase dimer structure (PDB ID: 7AJD) as a starting model by dividing the dimer into its rotary state 1 and rotary state 3 protomers, truncating all side chains and fitting these individually into the C2 tetramer cryo-EM map using the fit map function in UCSF ChimeraX and real space refinement in Coot(49). Dimers of IF1 were modeled by fitting the porcine tetramer IF1 models, mutating them to the sequence of bovine IF1, deleting the N-terminus up to glycine 23 and truncating all side chains. To construct the planar dimer model, the stator domain structure of bovine ATP synthase (PDB ID: 6ZIU), the F1c8-peripheral stalk structure (PDB ID: 6Z1U) and the bovine IF1 X-ray crystal structure (PDB ID: 1GMJ) were employed as structural references, truncated to their main chain and fitted into the planar dimer cryo-EM map. Both subunit 6.8PL and DAPIT were deleted from the model. The model of the IF1 bound rotary state 1 monomer F1 domain was constructed by employing the F1-peripheral stalk structure (PDB ID: 6YY0) as a starting model. After fitting and real space refinement using UCSF ChimeraX and Coot peripheral stalk subunits d, b, F6 and OSCP were truncated to their main chain and for subunit b its N-terminus was deleted up to aspartate 109, whereas for subunit d its C-terminus was deleted up to glutamine 112.

### Molecular Dynamics Simulations

A symmetric dimer model of bovine IF1 was prepared from the previously determined crystal structure(27) (PDBID: 1GMJ) with PyMOL by aligning the structure of chain A to that of chain B and by replacing the structure of chain B with the aligned structure chain A. The symmetric dimer model was then immersed in a cubic box of water with a side length of 158 Å using the Solution Builder(50) function of the CHARMM-GUI server(50). The N- and C-termini were capped with acetyl and *N*-methyl groups, respectively, and Lys49 of the 1GMJ structure was replaced by the original His residue. Potassium and chloride ions were added at a concentration of 0.15 M. The TIP3P model(51) was used for water molecules, the Amber ff14SB(52) force field was used for protein. His49 and His70 of each chain were protonated based on the comparison of the stability of the IF1 dimer interface between different protonation states in preliminary MD simulations.

After the energy of the system was minimized, the system was equilibrated in nine steps. Firstly, the system was gradually heated to 300 K in a 250-ps constant-NVT MD simulation with position restraints imposed on non-hydrogen atoms with a force constant of 10 kcal mol_−1_ Å_−2_. Then, the system was equilibrated in the NPT ensemble in the subsequent eight steps, in which MD simulations were conducted for 3 ns in total and the force constant of the position restraints was gradually reduced to zero. Finally, a production run was performed for 500 ns in the NPT ensemble without any restraints. The Monte Carlo barostat(53) was used to keep pressure at 1 bar, and the Bussi thermostat(54) was used to keep temperature at 300 K. Van der Waals interactions were calculated with a cutoff radius of 10 Å. The particle mesh Ewald (PME) method(55) was used to calculate electrostatic interactions. The SHAKE algorithm(56) was used to constrain the bond lengths involving hydrogen atoms to allow the use of a large (2 fs) time step. All the simulations were performed with the PMEMD module of Amber22(57). Trajectory analyses were conducted with the CPPTRAJ module of Amber22.

*Sus scrofa* and *Homo sapiens* models were generated from the symmetric dimer model of bovine IF1 by replacing bovine residues with *Sus scrofa* or *Homo sapiens* residues based on the BLAST sequence alignments(58). Specifically, R25, E29, K39, and A57 were replaced with K, D, R, and V, respectively, in the *Sus scrofa* model and R37, A38, K39, N51, S54, A57 were replaced with Q, S, R, E, V, and K, respectively, in the *Homo sapiens* model. Preparation of the solvated models and subsequent MD simulations were performed in a similar manner as described above.

The PTM model was generated by replacing Lys78 of the symmetric dimer model of bovine IF1 with N6-succinyl lysine (succK). The topology and the force field parameters of succK were generated by the residuegen module of Amber22. Specifically, a capped dipeptide, in which the α-amino and the α-carboxy groups of succK were capped with acetyl and *N*-methyl groups, respectively, was constructed and its geometry was optimized at the B3LYP/6-31G(d) level. The partial charges were calculated with the restrained electrostatic potential (RESP) method using the electrostatic potentials calculated at the HF/6-31G(d) level for the optimized structure. The quantum mechanical calculations were performed with Gaussian 16, revision C.02(59). The model was solvated using the LEaP module of Amber22. Potassium and chloride ions were placed around the protein with the SPLIT method(60) at a concentration of 0.15 M. The MD simulations were conducted under the same conditions as described above.

## Acknowledgements

We thank Tomitake Tsukihara and Yukio Morimoto for help at the initial stages of the project and Yuko Misumi for assistance with negative stain electron microscopy. We also thank Amelie Funakoshi and Bernhard C. Ludewig for graphical support..

## Author contributions

C.J., K.Y. and C.G. initiated the project; C.J. established and performed ATP synthase purification with assistance from G.K.; A.N. performed cryo-EM and image processing with assistance and supervision by K.M.; E.Y. and C.G. built models; D.S., C.G. and T.T. initiated MDS of IF1; Y.T. and T.T. designed, performed and interpreted MDS of IF1; T.T., D.S., K.Y. and C.G. wrote the manuscript with support from all other authors.

## Funding

We acknowledge funding from JSPS grant 20J40167 and 19K15749 (C. J.); the Naito Foundation Subsidy for Female Researchers after Maternity Leave (C. J.) and the Research Support Project for Life Science and Drug Discovery (Basis for Supporting Innovative Drug Discovery and Life Science Research (BINDS)) from AMED under Grant Number JP25ama121001 (to Masaki Yamamoto, G.K. and C.G.), JP25ama121027 (T.T.) and JP25am0101001 (D.S.).

## Data availability

Single-particle cryo-EM maps of bovine ATP synthase in its oligomeric forms of planar dimer, tetramer and monomer have been deposited at the Electron Microscopy Data Bank (EMDB); atomic models have been deposited at the protein data bank (PDB). Planar dimer: EMD-65237 (PDB 9VPB); tetramer, C2 symmetry imposed: EMD-65238 (PDB 9VPC); state 1 monomer, F1 focused: EMD-65239 (PDB 9VPD); tetramer: EMD-65240; tetramer, Fo focused map of B-state1: EMD-65241; state 2 monomer: EMD-65242; state 2 monomer, F1 focused map: EMD-65243; state 2 monomer, Fo focused map: EMD-65244; state 3 monomer: EMD-65245; state 3 monomer, F1 focused map: EMD-65246; state 3 monomer, Fo focused map: EMD-65247; state 1 monomer: EMD-65248; state 1 monomer, Fo focused map: EMD-65249. Raw cryo-EM image data sets are available at the Electron Microscopy Public Image Archive (EMPIAR), accession code EMPIAR-xxxxx, EMPIAR-xxxxx and EMPIAR-xxxxx.

## Notes

### Competing Interest Statement

The authors have declared no competing interest.

