## SupplementalMaterial for "A planar dimer of bovine ATP synthase"

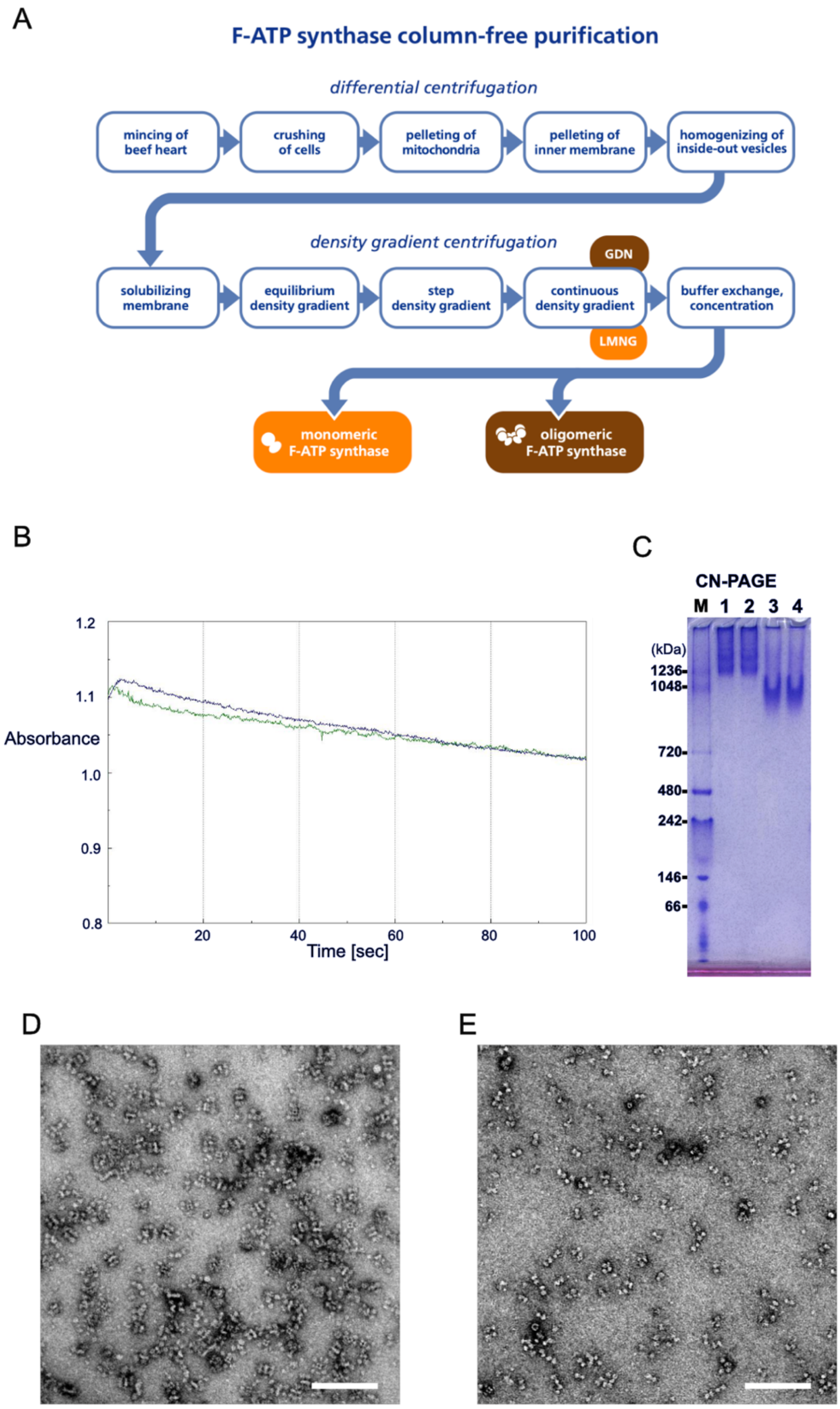


**Supplementary Figure 1:** Purification of oligomeric and monomeric bovine F-ATP synthase from bovine heart muscle tissue(17). (A) Workflow scheme. (B) ATP hydrolysis activity measurement of final oligomer fractions (n=2) indicating a very low ATP hydrolysis activity, which is in line with a high level of IF1 binding to all ATP synthase complexes of the preparation. (C) Clear native PAGE of final oligomer fractions (Lane 1, 2) and monomer fractions (Lane 3, 4), directly after purification (Lane 1&3) and after three days storage at

4 °C (Lane 2&4). Negative stain EM of (D) final oligomer fractions and (E) monomer fractions. Scale bar 100 nm.

| | **Cryo-EM structure of IF1 bound bovine ATP synthase planar dimer** | **Cryo-EM structure of IF1 bound bovine ATP synthase tetramer, C2 symmetry** | **Cryo-EM structure of IF1 bound bovine ATP synthase State 1 monomer F1 domain** |
|---|---|---|---|
| **PDB** | **9VPB** | **7AJD** | **9VPD** |
| **EMDB** | **EMD-65237** | **EMD-65238** | **EMD-65239** |
| **Data collection and processing** | | | |
| Magnification | 88,000 | | |
| Voltage (kV) | 300 | | |
| Electron exposure ($e^-/Å^2$) | 50 | | |
| Defocus range (µm) | 0.8 - 2.0 | | |
| Pixel size (Å) | 0.88 | | |
| Symmetry imposed | C1 | C2 | C1 |
| Initial particle images (no.) | 2,868,096 | 5,743,709 | 5,605,840 |
| Final particle images (no.) | 22,478 | 39,764 | 398,620 |
| Map resolution (Å) | 5.0 | 7.2 | 2.4 |
| FSC threshold | 0.143 | | |
| **Refinement** | | | |
| Initial model used (PDB code) | 6ZIU, 6Z1U, 1GMJ | 7AJD | 6YY0 |
| Model resolution (Å) | 8.0 | | 3.6 |
| FSC threshold | 0.5 | | |
| **Model composition** | | | |
| Nonhydrogen atoms | 49,950 | 101,921 | 27,808 |
| Protein residues | 10,166 | 20,620 | 3,844 |
| Ligands | 0 | 0 | 5MG, 3ATP, 3ADP |
| **R.m.s. deviations** | | | |
| Bond length (Å) | 0.006 | 0.005 | 0.007 |
| Bond angles (°) | 1.299 | 1.119 | 1.088 |
| **Validation** | | | |
| MolProbity score | 2.18 | 1.37 | 1.92 |
| Clash score | 14 | 1.9 | 7.63 |
| Rotamer outlier (%) | 0 | 0 | 0 |
| CaBLAM outliers (%) | 2.49 | 2.61 | 3.65 |
| **Ramachandran plot** | | | |
| Favored (%) | 90.72 | 93.64 | 91.5 |
| Allowed (%) | 8.20 | 6.26 | 8.5 |
| Disallowed (%) | 1.07 | 0.1 | 0 |

**Table S1 | Statistics of structural analysis:** map/model of planar dimer, tetramer and State 1 monomer F1 domain

| | Cryo-EM structure of IF1 bound bovine ATP synthase tetramer, C1 | Cryo-EM structure of IF1 bound bovine ATP synthase tetramer focused map B-state1 Fo | Cryo-EM structure of IF1 bound bovine ATP synthase State 2 monomer overall | Cryo-EM structure of IF1 bound bovine ATP synthase State 2 monomer focused F1 |
|---|---|---|---|---|
| EMDB | EMD-65240 | EMD-65241 | EMD-65242 | EMD-65243 |
| **Data collection and processing** | | | | |
| Magnification | 88,000 | | | |
| Voltage (kV) | 300 | | | |
| Electron exposure ($e^-$/Å$^2$) | 50 | | | |
| Defocus range (μm) | 0.8 - 2.0 | | | |
| Pixel size (Å) | 0.88 | | | |
| Symmetry imposed | C1 | C1 | C1 | C1 |
| Initial particle images (no.) | 5,743,709 | 5,743,709 | 5,605,840 | 5,605,840 |
| Final particle images (no.) | 20,949 | 128,686 | 117,760 | 117,760 |
| Map resolution (Å) | 7.9 | 4.0 | 2.6 | 2.6 |
| FSC threshold | 0.143 | | | |

| | Cryo-EM structure of IF1 bound bovine ATP synthase State 2 monomer focused Fo | Cryo-EM structure of IF1 bound bovine ATP synthase State 3 monomer overall | Cryo-EM structure of IF1 bound bovine ATP synthase State 3 monomer focused F1 | Cryo-EM structure of IF1 bound bovine ATP synthase State 3 monomer focused Fo |
|---|---|---|---|---|
| EMDB | EMD-65244 | EMD-65245 | EMD-65246 | EMD-65247 |
| **Data collection and processing** | | | | |
| Magnification | 88,000 | | | |
| Voltage (kV) | 300 | | | |
| Electron exposure ($e^-$/Å$^2$) | 50 | | | |
| Defocus range (μm) | 0.8 - 2.0 | | | |
| Pixel size (Å) | 0.88 | | | |
| Symmetry imposed | C1 | C1 | C1 | C1 |
| Initial particle images (no.) | 5,743,709 | 5,743,709 | 5,743,709 | 5,743,709 |
| Final particle images (no.) | 398,620 | 488,235 | 488,23 | 36,118 |
| Map resolution (Å) | 3.6 | 2.4 | 2.2 | 3.3 |
| FSC threshold | 0.143 | | | |

| | Cryo-EM structure of IF1 bound bovine ATP synthase State 1 monomer overall | Cryo-EM structure of IF1 bound bovine ATP synthase State 1 monomer focused Fo |
|---|---|---|
| EMDB | EMD-65248 | EMD-65249 |
| **Data collection and processing** | | |
| Magnification | 88,000 | |
| Voltage (kV) | 300 | |
| Electron exposure ($e^-$/Å$^2$) | 50 | |
| Defocus range (μm) | 0.8 - 2.0 | |
| Pixel size (Å) | 0.88 | |
| Symmetry imposed | C1 | C1 |
| Initial particle images (no.) | 5,743,709 | 5,743,709 |
| Final particle images (no.) | 398,620 | 117, 760 |
| Map resolution (Å) | 2.4 | 3.1 |
| FSC threshold | 0.143 | 0.143 |

**Table S2 | Statistics of structural analysis:** additional maps

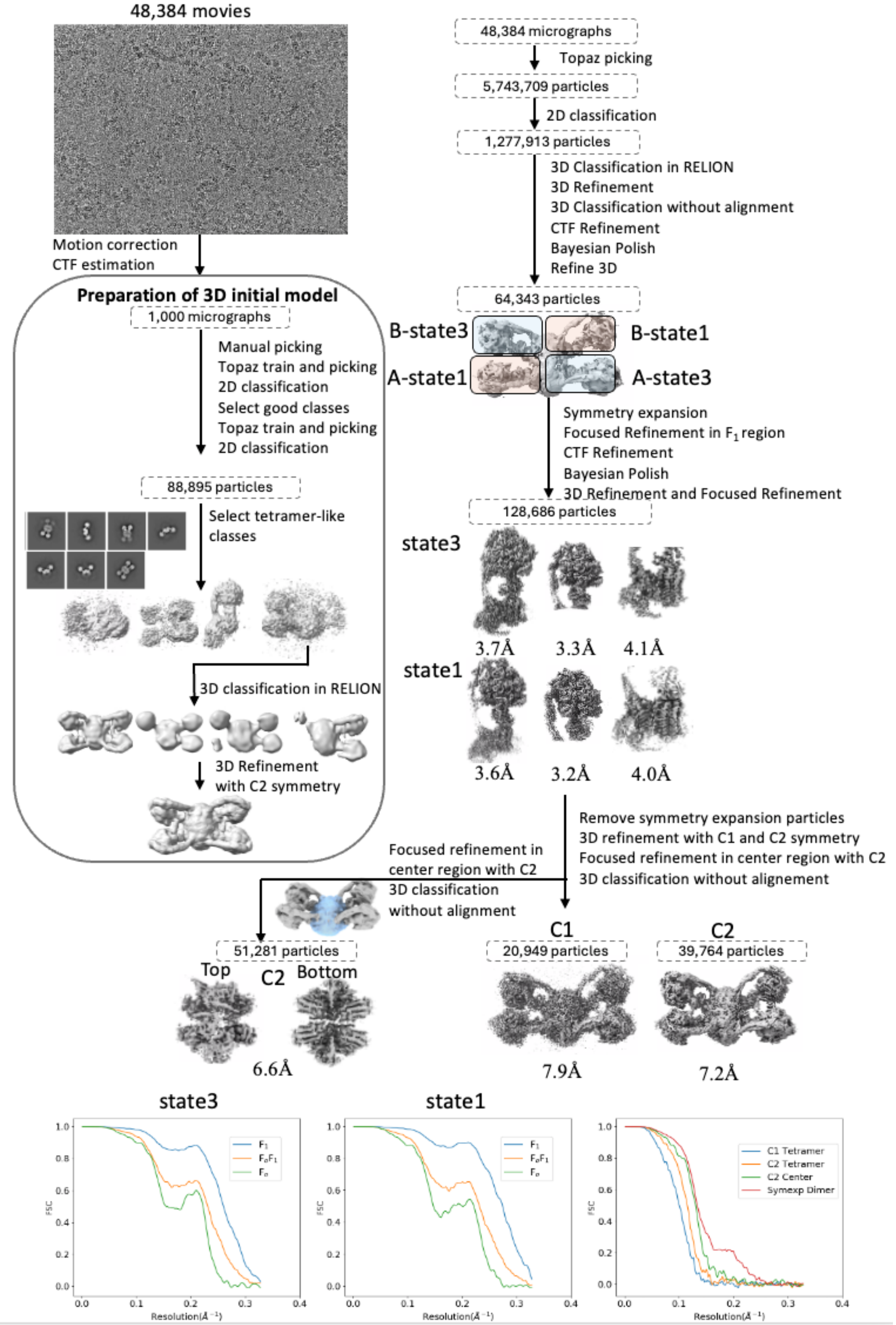


**Supplementary Figure 2A**: Image processing bovine tetramer

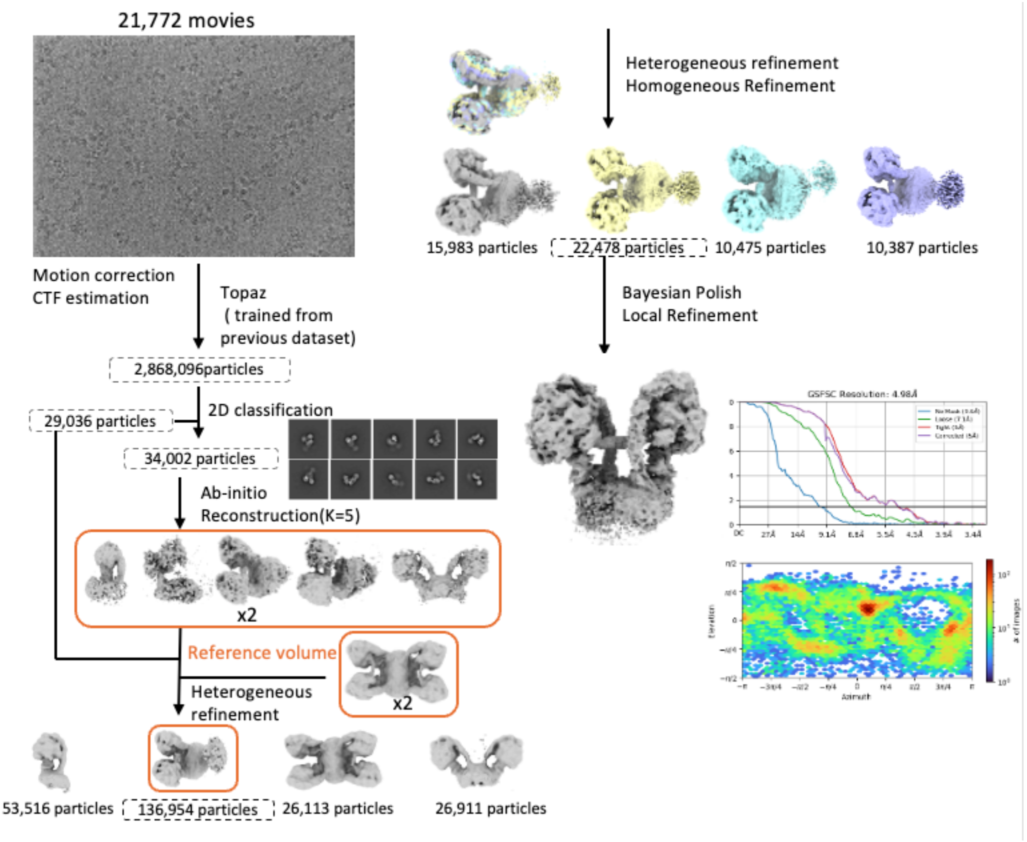


**Supplementary Figure 2B:** Image processing bovine planar dimer

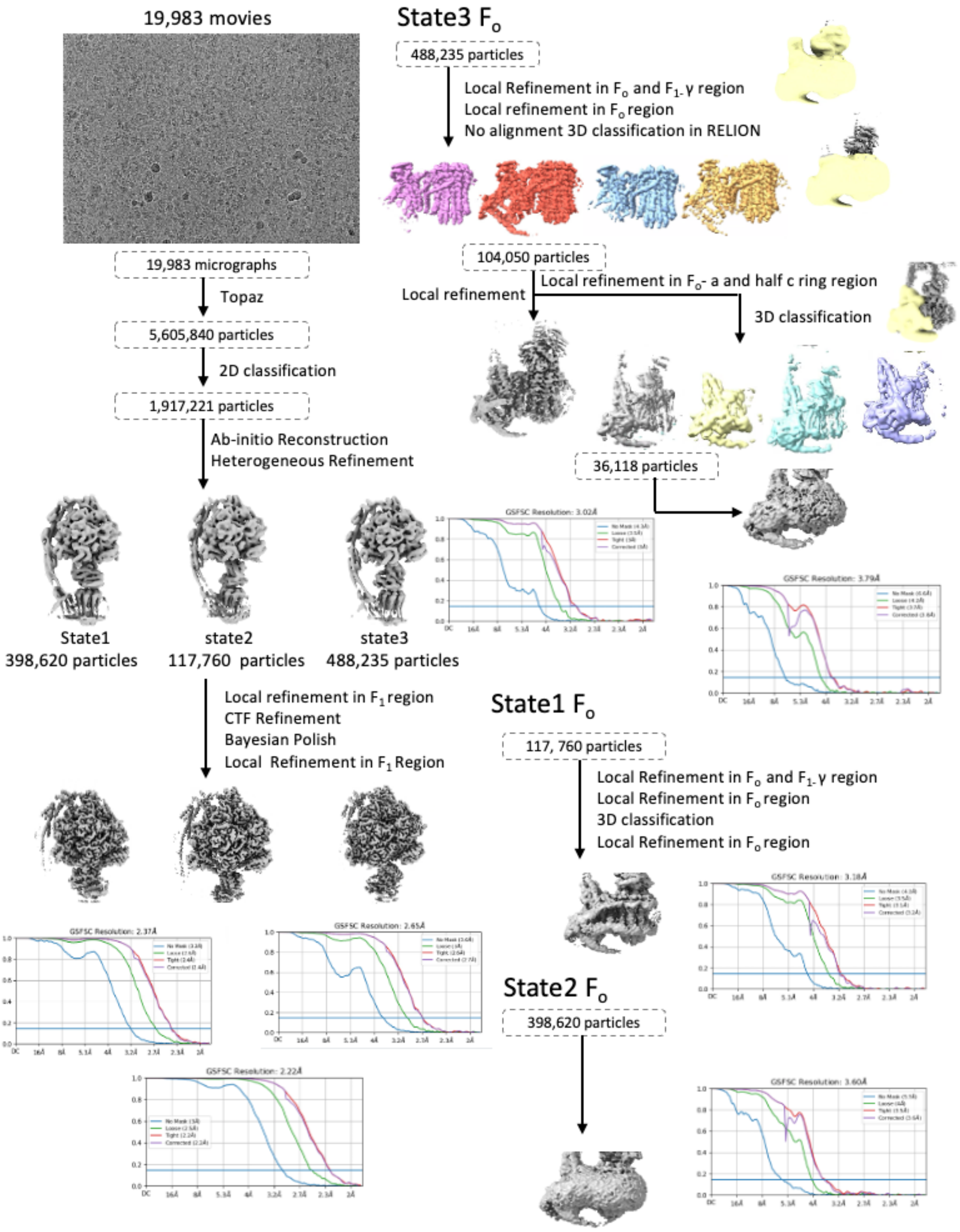


**Supplementary Figure 2C:** Image processing bovine monomer

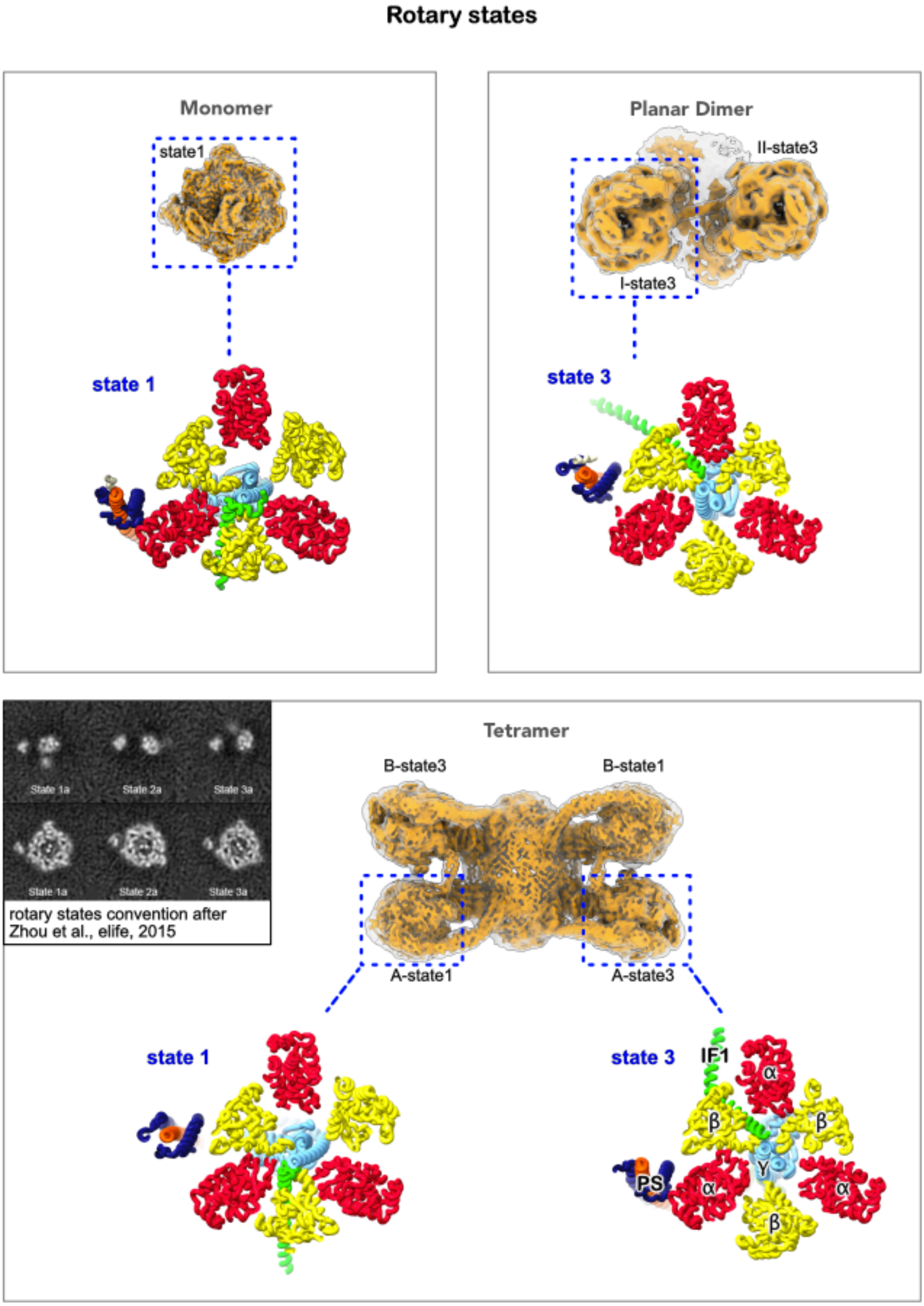


**Supplementary Figure 3:** Rotary states(21) of ATP synthase for the state 1 monomer (upper left panel), the planar dimer (upper right panel) and the tetramer (lower panel).

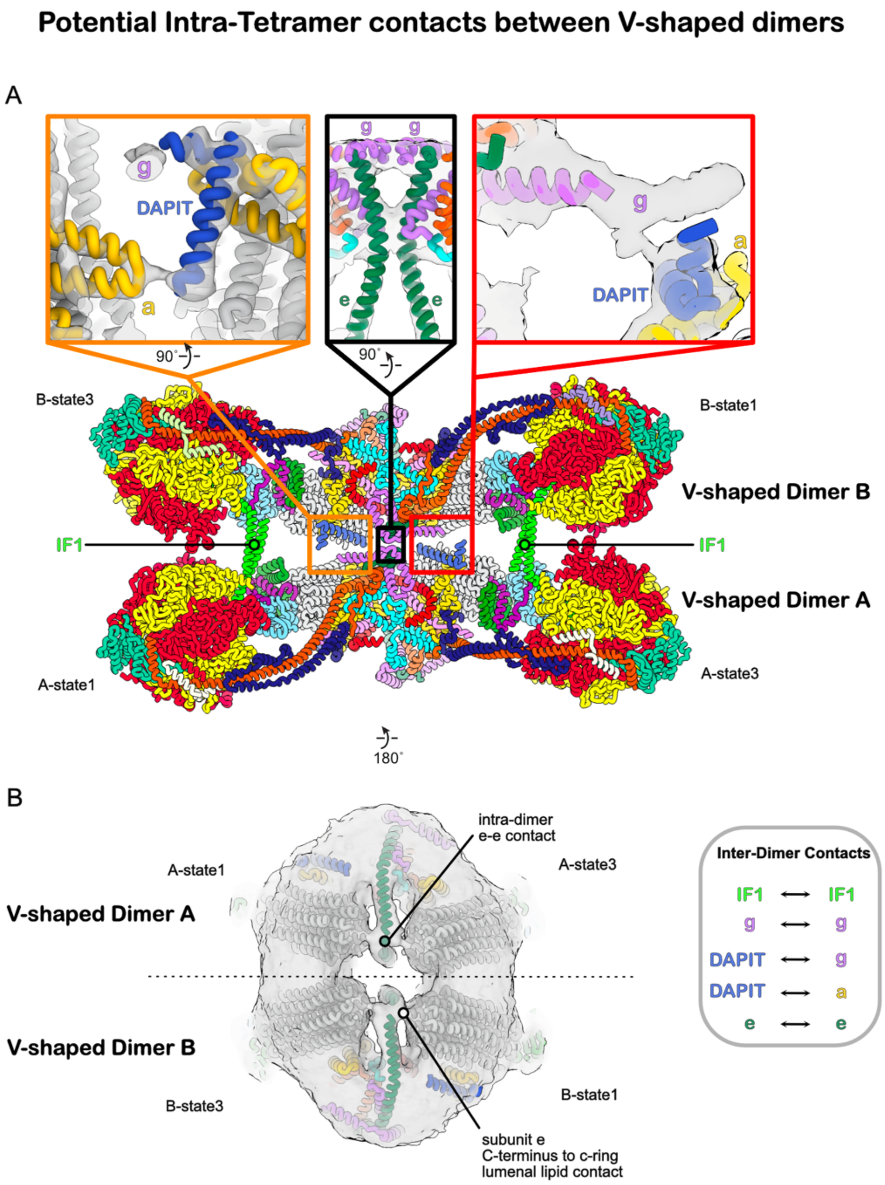


**Supplementary Figure 4:** Potential intra-tetramer contacts between V-shaped dimers for the IF1 bound bovine ATP synthase tetramer.

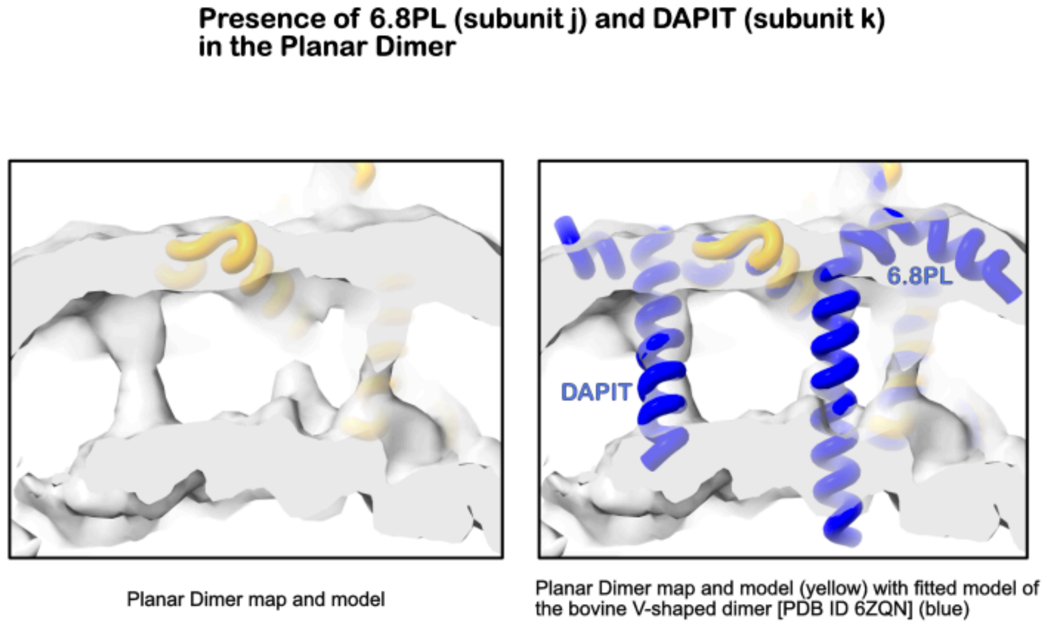


**Supplementary Figure 5: Presence vs absence of cryo-EM density for 6.8PL (subunit j) and DAPIT (subunit k) in the planar dimer.** The presence of DAPIT is supported by clear density indicative of a single transmembrane helix, whereas transmembrane density in the expected position of 6.8PL is absent, suggesting the absence of 6.8PL in the planar dimer.

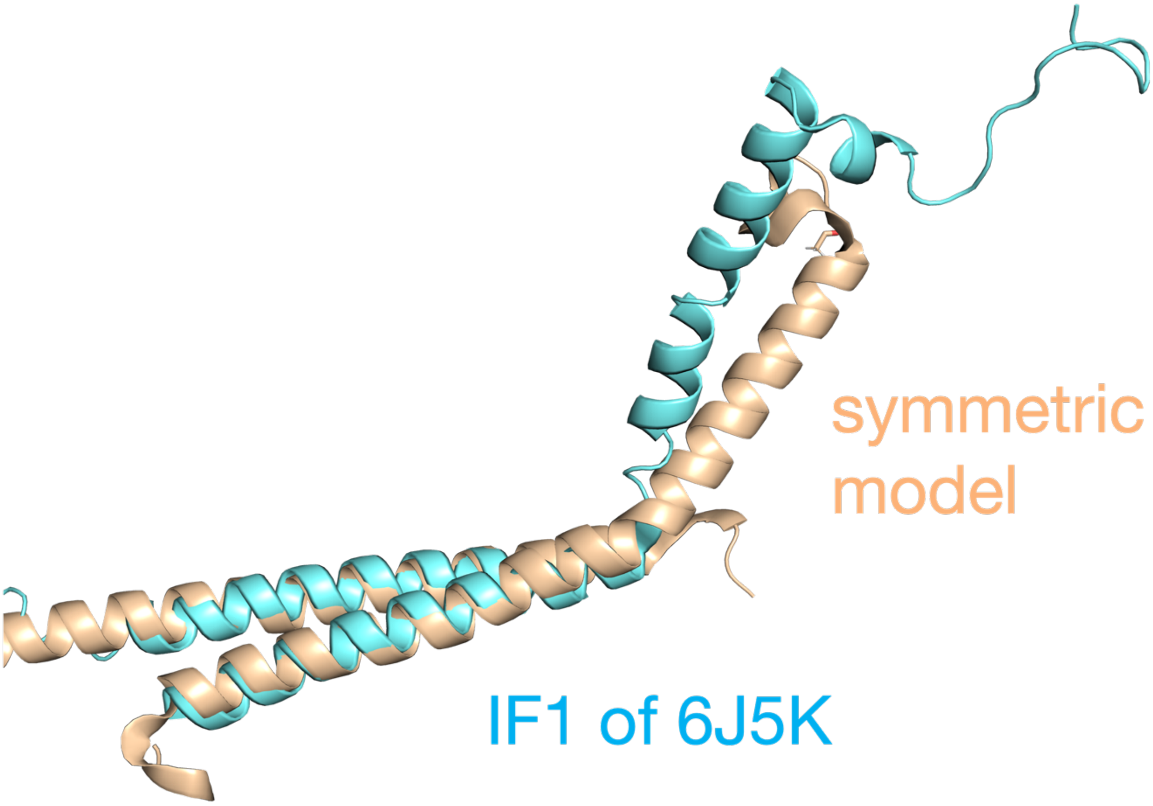


**Supplementary Figure 6:** Alignment of a snapshot from the MD simulation for the symmetric dimer model of bovine IF1 (brown) with the IF1 dimer structure from the cryo-EM structure of the ATP synthase tetramer bound with IF1 (blue, PDB ID: 6J5K).

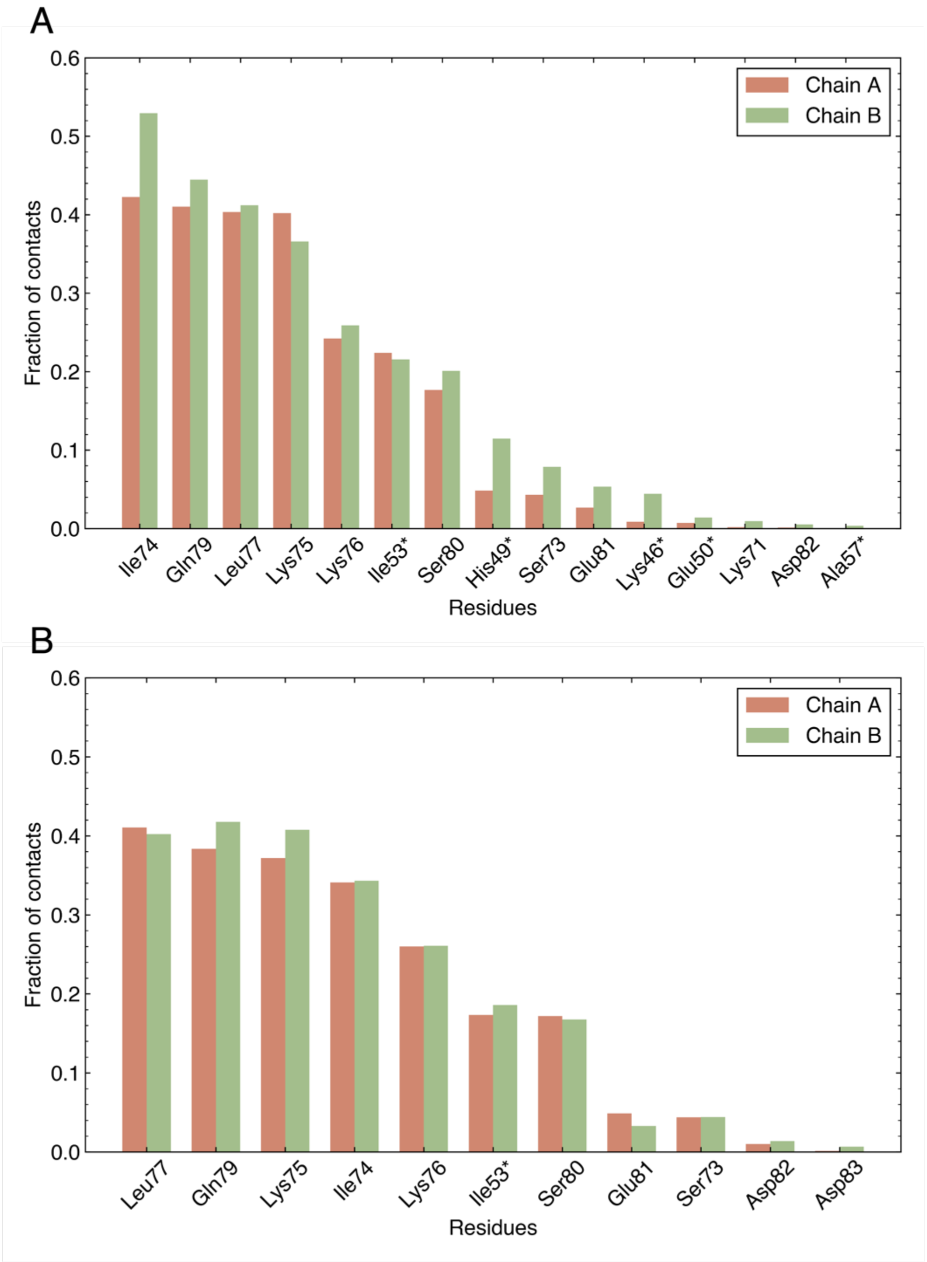


**Supplementary Figure 7:** Bar charts showing fractions of times when succinylated Lys78 (SuccK) (A) and original Lys78 (B) form contacts with neighboring residues from chains A (brown) and B (green) in the 500-ns MD simulations for the post-translationally modified and the original models of the bovine IF1 dimer, respectively. Asterisks indicate that the contacts are formed between residues from different chains.

**Supple. Movie md1**

Movie of the 500 ns MD simulation trajectory for the symmetric dimer model of bovine IF1. Protein backbone structures are shown in a ribbon representation.

**Supple. Movie md2**

Movie of the 500 ns MD simulation trajectory for the model with the post-translational modification. Protein backbone structures are shown in a ribbon representation. Succinylated Lys78 is shown in a stick representation.

**Supple. Movie md3**

Movie of the 500 ns MD simulation trajectory for porcine IF1 dimer model. Protein backbone structures are shown in a ribbon representation.

**Supple. Movie md4**

Movie of the 500 ns MD simulation trajectory for human IF1 dimer model. Protein backbone structures are shown in a ribbon representation.

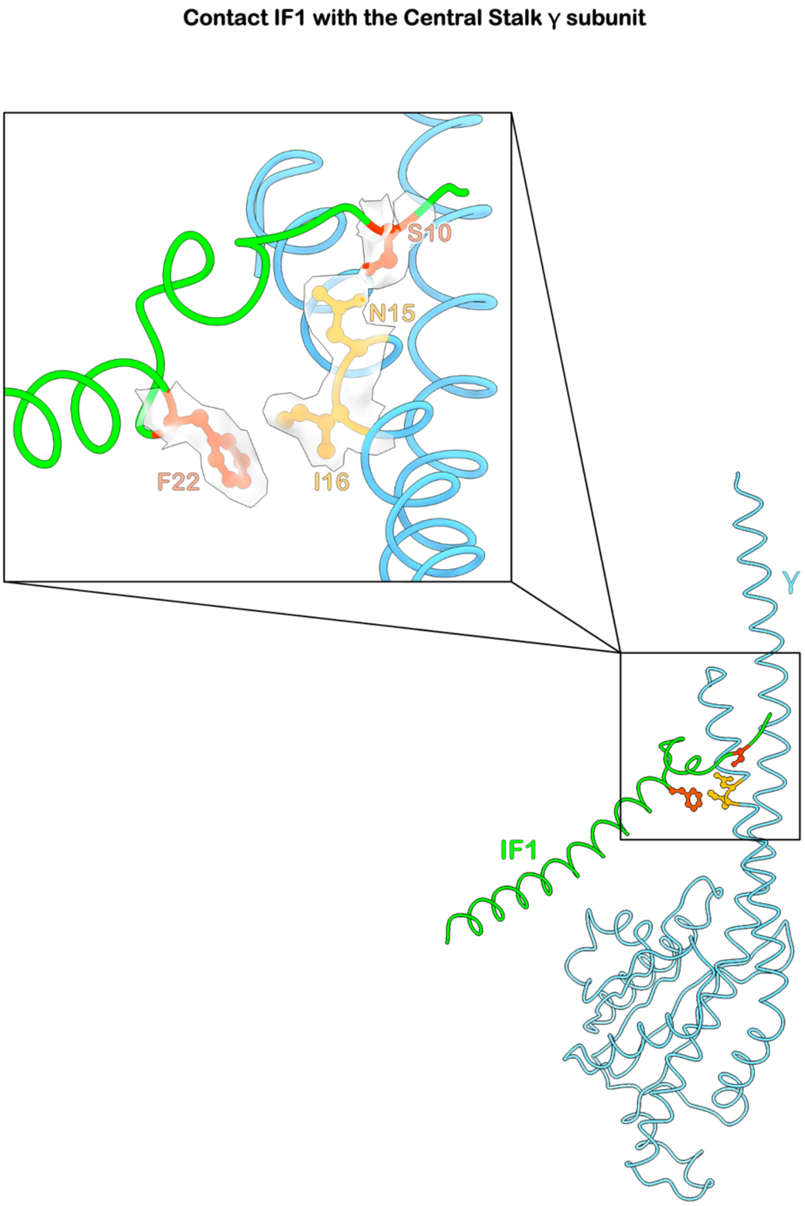


**Supplementary Figure 8:** Crucial contacts between the N-terminal domain of IF1 and the central stalk subunit γ that have been shown to be important for the directionality of IF1 inhibition of rotational catalysis are visualized in the intact bovine ATP synthase as expected from the IF1-fragment F1 X-ray crystal structure(31) (PDB ID 2v7q).

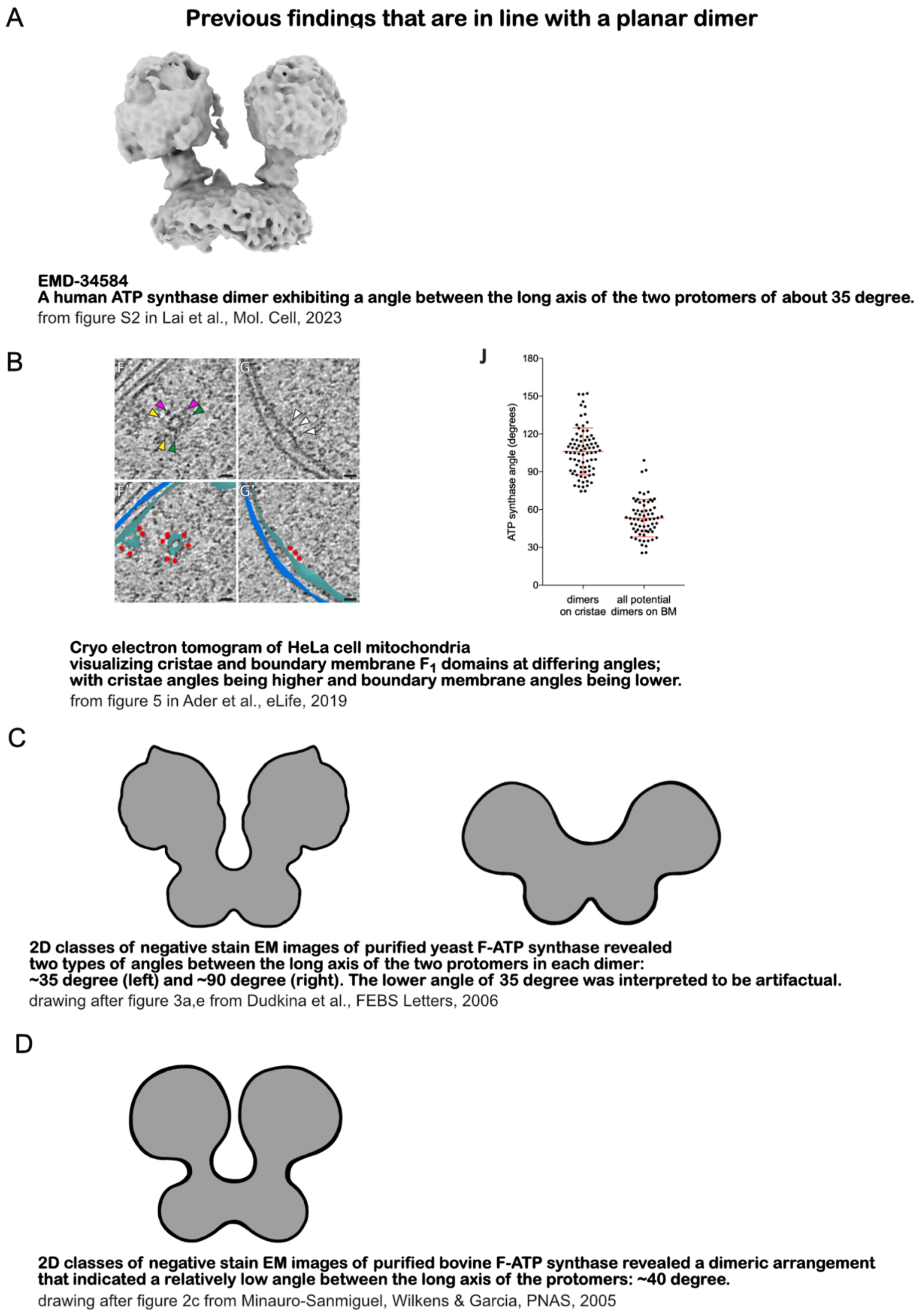


**Supplementary Figure 9:** Findings of previous reports that are in support of a planar dimer. Several reported data on mitochondrial ATP synthase from mammalian cells or yeast are in support of the existence of a planar dimer. Lack of strong structural data, however, rendered the existence and functional role of an oligomeric form different from the canonical V-shaped dimer a more speculative part of mitochondrial biology. The strongest indication for an alternative architecture of mammalian ATP synthase oligomers at the inner boundary membrane (IBM) stems from cryo electron tomography performed on HeLa cell mitochondria (see B).

**Supporting Discussion**

*In situ versus in vitro*

Since we aimed to investigate the oligomeric state of bovine ATP synthase from a natural source it might be asked, if *in situ* cryo-ET would not be the more suitable tool. In principle the answer would be affirmative; however, current technical limitations render the discovery of minor subpopulations of F-ATP synthase/hydrolase oligomeric states at relatively low copy number per mitochondrion very challenging. Of note, even cryo-ET specialized labs with a long and successful track record in the field still prefer to work on single cell model organisms for in situ structure investigation of mitochondrial F-ATP synthase/hydrolase. On the other hand, the structure of the planar dimer reported here could be used in future studies as a search template in cryo electron tomograms of mammalian mitochondria to pinpoint its exact mitochondrial location. In the long term, as advocated since early studies on the oligomeric state of mitochondrial ATP synthase, combining in situ and in vitro experiments will provide the most comprehensive insights into its oligomeric organization.

*Subunit assignment for the $F_o$ domain*

Subunit assignment of the $F_o$ domain of the porcine tetramer remains controversial(22). In particular, assigning the c-ring lumenal density to an alpha-helix belonging to the readily dissociable(23) supernumerary subunit 6.8PL (equivalent to subunit j in yeast) has been subject to criticism. In our bovine tetramer-focused $F_o$ map of the B-state1 protomer at 4.0 Å overall resolution, the density for the c-ring α-helices is well-defined, clearly visualizing the α-helical pitch and maintaining connectivity even at high threshold levels. In stark contrast, c-ring luminal density is almost absent at high thresholds, and stays completely disconnected in the center of the c-ring lumen even at threshold levels close to noise. This observation refutes the interpretation that a single polypeptide α-helix resides within the c-ring lumen of mammalian tetrameric ATP synthase(18), instead reinforcing the longstanding hypothesis that lipid molecules occupy the c-ring lumen(24, 25). What specific lipid molecules occupy the upper (matrix side) and lower (intercristae space side) leaflet positions cannot be determined on the basis of our or other available density maps. Nevertheless, due to their large size, cardiolipins can be ruled out and the suggested presence of a lysolipid in the lower position appears reasonable(26). In summary, we concur with the subunit assignment proposed by the Walker lab for bovine ATP synthase(22) and utilized PDB ID 6ZQN as our initial model. This subunit assignment aligns well with those proposed for ovine(26) and human ATP synthase monomers(27, 28) based on cryo-EM maps, as well as with predictions from AlphaFold3(29) for human and bovine ATP synthase monomers(30). As a result, the bovine tetramer structure described here provides strong evidence that the position of subunits within the $F_o$ domain do not undergo radical changes as the result of tetramer formation.

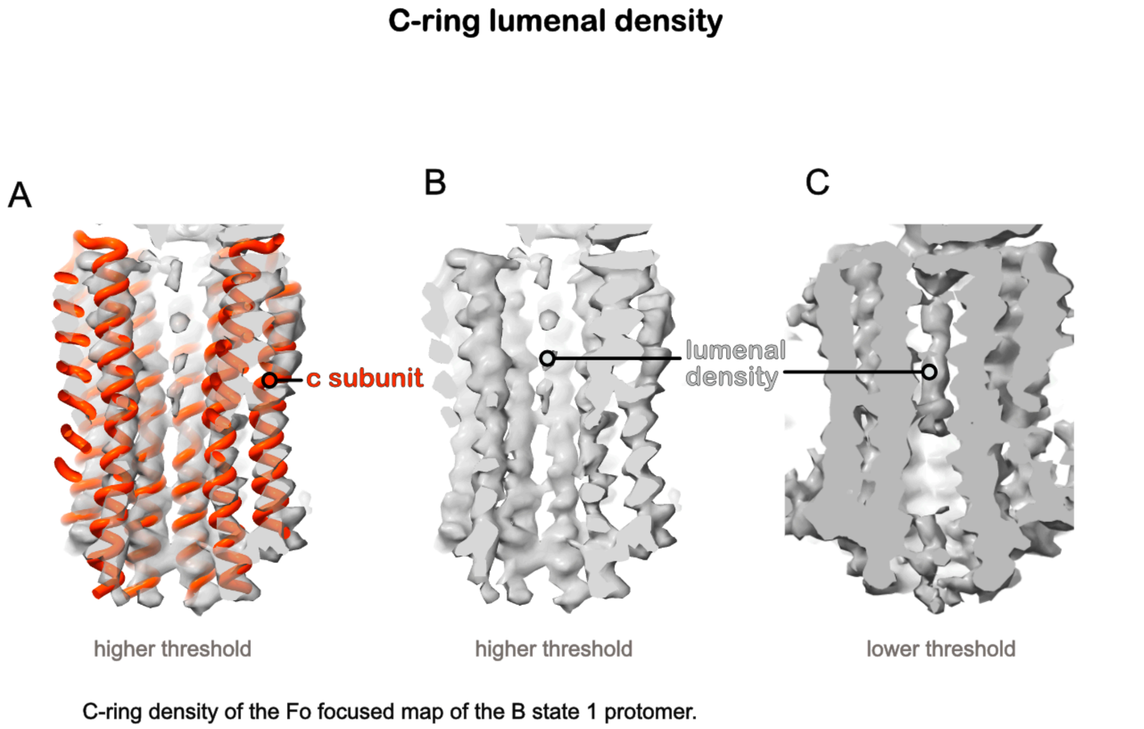


**Supplementary Figure 10:** Lumenal density of the c-ring as visualized for the Fo focused map from the bovine tetramer clearly indicates it to be not stemming from an alpha helical peptide.

*Possibility of artifactual complex formation by tetramer breakdown*

At low threshold our map of the planar dimer indicates the presence of a third, loosely attached ATP synthase complex. Therefore, it is prudent to ask, if the planar dimer might not be the artifactual result of a broken oligomer of V-shaped dimers. However, an IF1 bound planar dimer that is the artifactual result from IF1 bound tetramers would require the unbinding of the IF1 dimerization domain, monomerization of both V-shaped ATP synthase dimers, precise re-binding of the two state 3 ATP synthase monomers and finally detachment of the two $\alpha_{PS}$ from their respective peripheral stalks. Moreover, these events would have to produce sufficient numbers of complexes to allow structure determination at a level that is comparable to that of the ATP synthase tetramer. An overall highly unlikely scenario, which gives us confidence that the planar dimer existed in the inner mitochondrial membrane prior to solubilization (see also Supp.Fig.11&12).

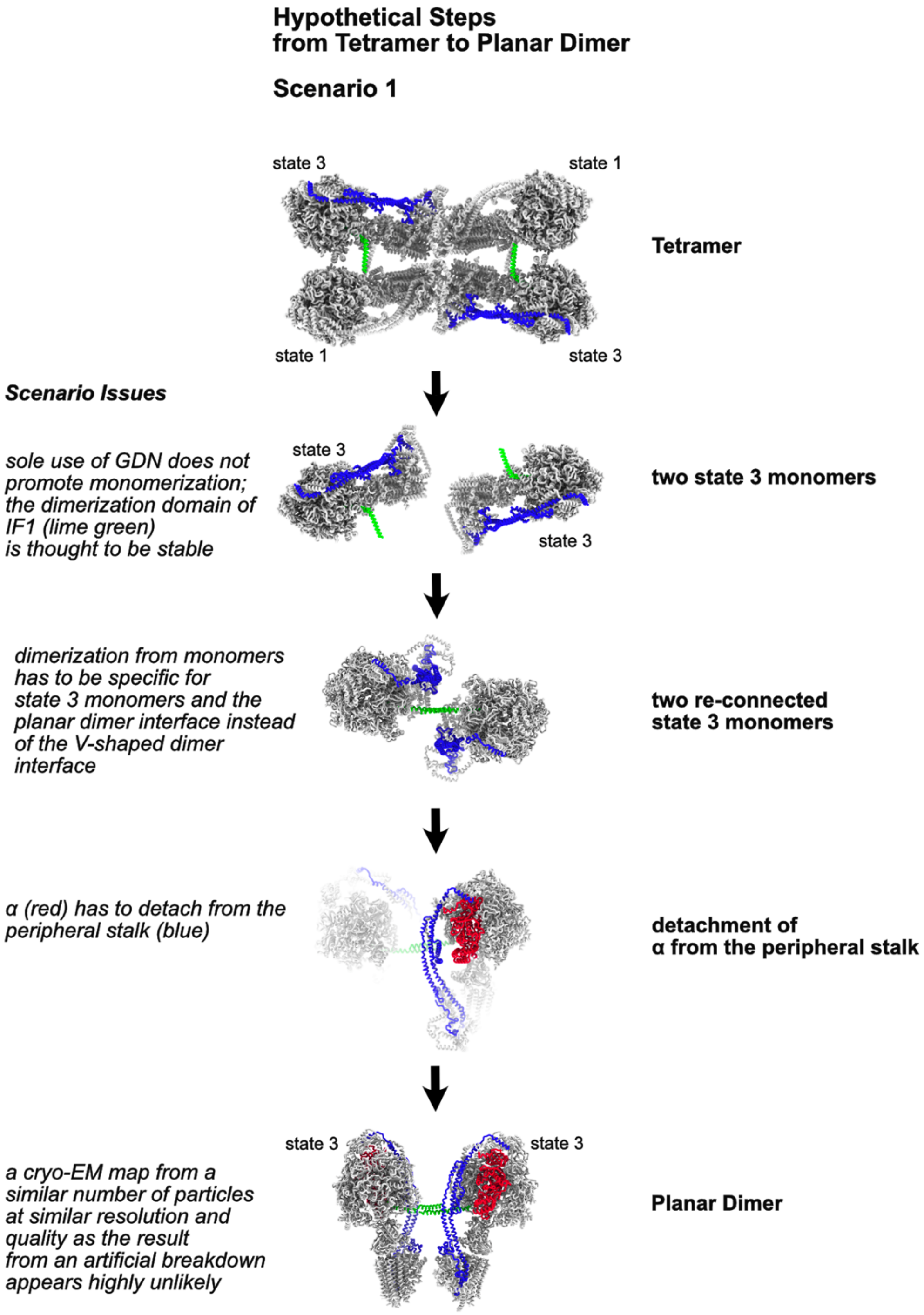


**Supplementary Figure 11:** A non-plausible scenario tetramer breakdown to planar dimer

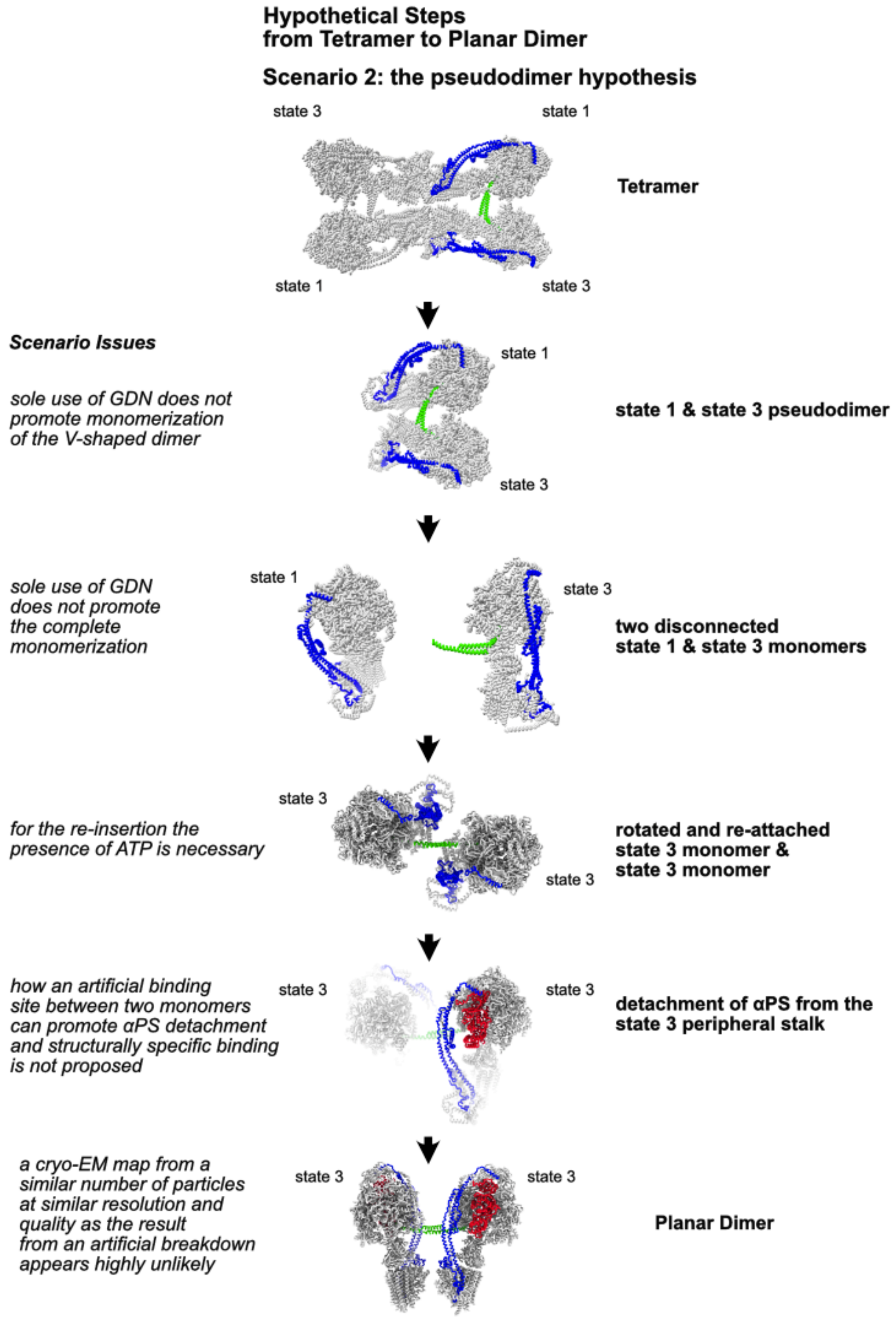


**Supplementary Figure 12:** A second non-plausible scenario tetramer breakdown to planar dimer based on the pseudo-dimer hypothesis from Dudkina et al., 2006
